# Acetoacetate suppresses colon cancer via an MR1-MAIT axis

**DOI:** 10.1101/2025.10.29.685375

**Authors:** Slater L. Clay, Geniver El Tekle, Edrees H. Rashan, Diogo Fonseca-Pereira, Yern-Hyerk Shin, Natalia Andreeva, Geicho Nakatsu, Ayesha F. Lobo, Sway P. Chen, Sena Bae, Monia Michaud, Jonathan N. Glickman, Jon Clardy, Matthew G. Vander Heiden, Wendy S. Garrett

## Abstract

Colorectal cancer (CRC) is a leading cause of cancer mortality and additional preventative, and therapeutic strategies are urgently needed. Ketogenic diets have mixed effects on tumorigenesis and compliance is challenging. Exogenous ketones, β-hydroxybutyrate (βHB) or acetoacetate (AcAc), offer an alternative approach. While βHB has been investigated, the anti-cancer effects of AcAc are poorly defined. Here, we show that orally administering ethyl AcAc (EAA) suppresses tumor growth in several pre-clinical CRC models. Single-cell RNA sequencing, flow cytometry, and genetic and antibody-mediated depletion studies reveal that EAA selectively expands and activates cytotoxic mucosal-associated invariant T (MAIT) cells in an MHC class I-related protein 1 (MR1)-dependent manner. EAA increases MR1 expression by tumor monocytes, which is recapitulated in human cell cultures, where AcAc and 5-amino-6-D-ribitylaminouracil (5-A-RU) induce MAIT cell expansion and tumor killing. Mechanistically, AcAc converts to methylglyoxal, combining with microbially-derived 5-A-RU to generate 5-(2-oxopropylideneamino)-6-D-ribitylaminouracil (5-OP-RU), a potent MR1 ligand. These findings identify an AcAc-MR1-MAIT cell axis as a potential immunotherapy approach for CRC therapy.

## MAIN

Colorectal cancer (CRC) is a highly prevalent malignancy and a leading cause of cancer-related mortality worldwide^1,2^. Compliance with screening is challenging, prognosis for advanced disease is poor, and early-onset CRC is increasing, all highlighting the urgent need for new preventive and therapeutic strategies^3,4^. Ketogenic diets, which promote endogenous ketone production, have gained attention for their effects on intestinal immunity and tumorigenesis^5–8^. However, adherence to these restrictive diets can be difficult, and they exert a wide range of potentially harmful physiological effects^9–11^. To delineate the role of ketones from other dietary effects, there is increasing interest in administration of exogenous ketones as a therapeutic approach^5,12–15^. β-hydroxybutyrate (βHB) and acetoacetate (AcAc) are the most abundant ketones; βHB has received the most attention and acetoacetate (AcAc) is understudied^16,17^, due in part to its instability, which could limit its utility as a therapeutic.

To evaluate the effects of exogenous AcAc on colonic tumorigenesis, we used an orally-delivered esterified form of AcAc in several models of CRC. This approach increased AcAc levels in the serum and in tumors and reduced tumor burden and neoplastic progression in both genetically engineered (GEMM) and orthotopic mouse CRC models. Notably, AcAc treatment increased intra-tumoral mucosal-associated invariant T (MAIT) cells. MAIT cells are polyfunctional effectors that contribute to antimicrobial defense, inflammation and tissue repair^18–21^. MAIT cells can be rapidly activated by inflammatory signals and small molecule ligands presented by the MHC class I-like molecule MR1^22–24^. The best-characterized MR1 ligand, 5-(2-oxopropylideneamino)-6-D-ribitylaminouracil (5-OP-RU), is generated from the bacterially derived riboflavin intermediate 5-amino-6-D-ribitylaminouracil (5-A-RU) and methylglyoxal (MGO)^25–27^.

While MAIT cells affect intestinal immunity and alter tumor progression, both pro- and anti-tumor effects have been reported^28–34^. Here, we show that AcAc generates 5-OP-RU, a potent MAIT cell-activating MR1 ligand, in the presence of 5-A-RU. This increases and activates anti-tumor MAIT cells in mouse pre-clinical models and human cell cultures. Genetic and antibody-mediated MR1 depletion abrogated the anti-tumor effects of AcAc. Collectively, our results reveal a previously unrecognized link between AcAc, MR1 antigen presentation, and anti-tumor MAIT cells, highlighting the potential of targeting this pathway as a therapeutic strategy.

### Oral EAA increases in vivo AcAc and inhibits tumor growth in multiple CRC models

To circumvent the instability of AcAc, we administered ethyl AcAc (EAA) in drinking water, which is hydrolyzed to AcAc in vivo (Extended Data Fig. 1a). EAA supplemented drinking water increased serum AcAc but not βHB (Fig. 1a,b), enabling non-invasive delivery of AcAc suitable for CRC models in which tumors develop over many weeks to months (Extended Data Fig. 1b-d). EAA-treated mice showed elevated AcAc levels (0.33 mM [0.15–0.51]) (Fig. 1b) that parallel those observed in humans undergoing nutritional ketosis^15,35^. EAA provided ad libitum for 3 weeks did not affect drinking water intake, body weight, or serum glucose levels (Extended Data Fig. 1e-g).

**Figure 1.**
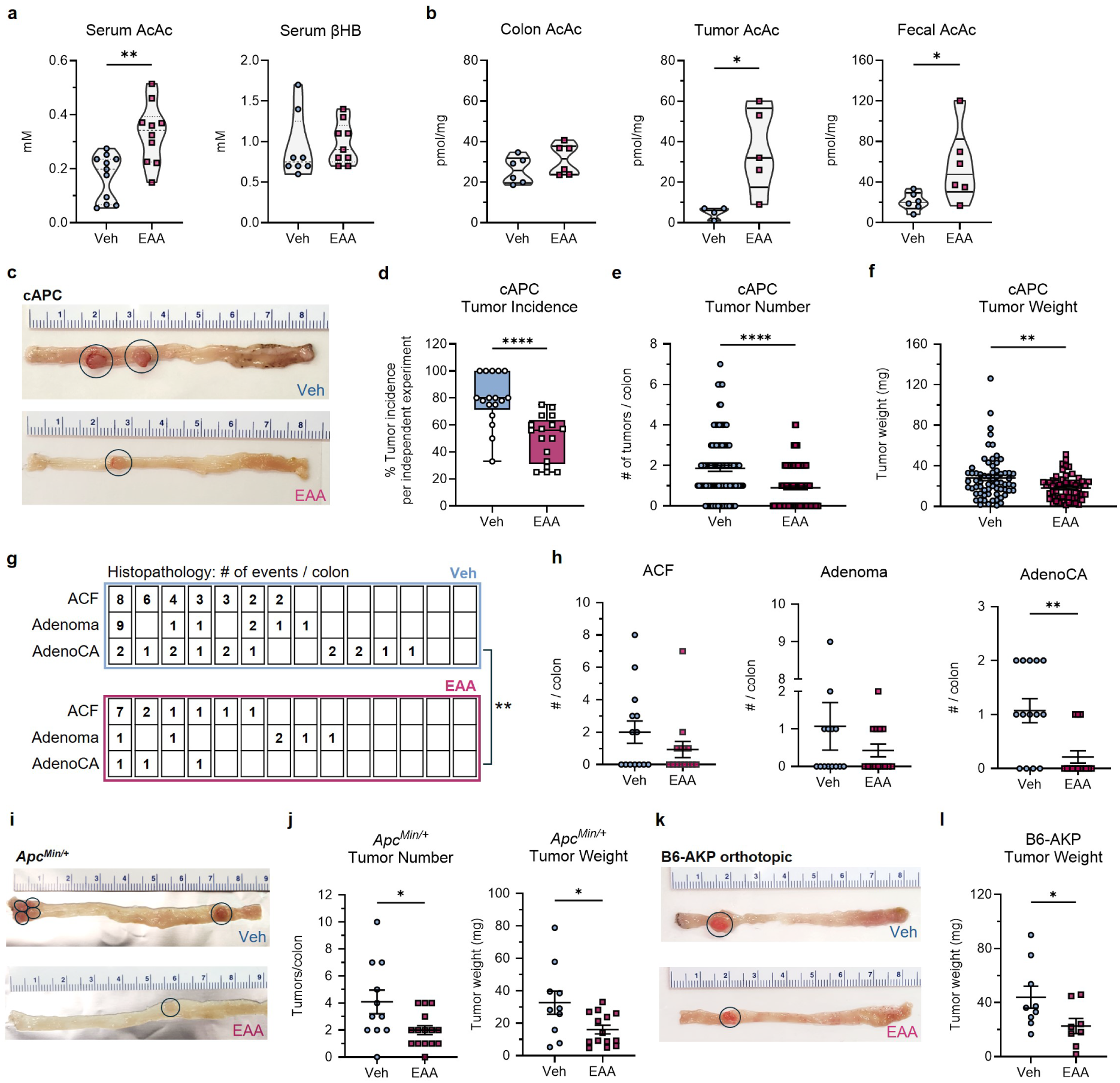
Oral ethyl acetoacetate (EAA) inhibits tumor growth. **a**, Serum concentrations of AcAc and β-hydroxybutyrate (βHB) in *Cdx2*^Cre^ x *Apc*^flox/+^ (cAPC) mice following treatment. **b**, AcAc levels in tumor, adjacent colon, and fecal samples. **c**, Representative images of colon tumors in cAPC mice (left, distal; right, proximal) with visible tumors circled. **d**, Tumor incidence in 17 independent cAPC experiments. Each data point represents tumor incidence of an independent cohort. **e**, Tumor number per colon (n = 114-125 / group) and **f**, tumor weight (n = 56-66 / group). **g**, Lesion grade and number assessed by histopathology. Columns represent individual animals (n = 14 / group), ordered by number of events; rows indicate lesion grade: aberrant crypt foci (ACF), adenoma, or adenocarcinoma (AdenoCA). **h**, Number of lesions of each grade by treatment group. **i**, Representative images of colon tumors in *Apc*^Min/+^ mice (left, distal; right, proximal) with visible tumors circled. **j**, *Apc*^Min/+^ tumor numbers and weight following treatment (n = 10-15 / group). **k**, Representative images of orthotopic AKP tumors in C57BL/6 (B6) mice (left, distal; right, proximal) with visible tumors circled. **l**, B6-AKP tumor weight following treatment (n = 8-9 / group). Data are representative of two independent experiments (**a**,**b**), 17 independent experiments (**c**-**f**), three independent experiments (**g**-**j**), and four independent experiments (**k**,**l**). Lines and error bars represent mean ± s.e.m. Statistical significance was determined using the Mann-Whitney *U* test (**a**,**b**,**e**-**h**,**j**,**l**) or two-tailed Wilcoxon matched-pairs signed-rank test (**e**). \**P* < 0.05, \*\**P* < 0.01, \*\*\*\**P* < 0.0001.

To evaluate the potential benefits or risks of EAA supplementation in the context of colorectal tumorigenesis, we first tested EAA in *Cdx2*^Cre^ x *Apc*^flox/+^ (cAPC) mice, which harbor an *Apc* deletion in large intestinal epithelial cells and develop microsatellite stable colonic tumors sporadically with age, as seen in human disease^36,37^. EAA was administered from 6-8 weeks of age until 18-20 weeks (Extended Data Fig. 1b), coinciding with tumor presence but preceding marked health deterioration. In addition to increasing serum AcAc (Fig. 1a), EAA elevated AcAc levels in tumors and fecal samples (Fig. 1c). Strikingly, EAA treatment consistently reduced tumor burden as assessed by incidence, number, and weight (Fig. 1d-g), in both sexes (Extended Data Fig. 1h), and blunted neoplastic progression from aberrant crypt foci or adenoma to adenocarcinoma (Fig. 1h,i).

To evaluate EAA’s potential as a therapeutic, in contrast with its effectiveness as a preventative agent (Extended Data Fig. 1b-d), we initiated EAA delivery at 14-16 weeks of age, the timeframe when cAPC mice have visible colonic tumors (Extended Data Fig. 1i-k). EAA decreased tumor number and weight, supporting efficacy against pre-existing tumors (Extended Data Fig. 1l-m). To test if anti-tumor effects were AcAc-specific or a general effect of ketones, we administered an esterified βHB (EβHB: Extended Data Fig. 1n), which increased serum βHB but did not reduce tumor burden (Extended Data Fig. 1o-q), indicating an anti-tumor effect specific to exogenous AcAc.

Next, we tested EAA in two additional CRC models: *Apc^Min/+^* mice treated with the mucosal disruptant dextran sulfate sodium (DSS) which induces inflammation and accelerates colonic tumorigenesis (Extended Data Fig. 1c)^38^, and an orthotopic CRC model which employs *Apc*-null*, Kras^G12D^ and Trp53*-null (AKP) tumor colonoids (Extended Data Fig. 1d)^39^. In both, EAA reduced tumor number and mass (Fig. 1j-m). These data show that oral EAA administration reduces colonic tumor burden across several distinct preclinical CRC models.

### EAA increases intra-tumoral MAIT cell abundance and anti-tumor effector function

To identify changes underlying the reduced tumor burden with EAA, we performed single cell RNA sequencing (scRNA-seq) on cAPC tumors. Transcriptomic analysis revealed high immune infiltration with distinct myeloid, lymphocyte, epithelial, tumor, and stromal clusters (Fig. 2a; Supplementary Table 1). Although EAA did not alter the abundance of major cell populations (Extended Data Fig. 2a), an analysis of the lymphocyte cluster containing T cells, NK cells, and ILCs (cluster 2) revealed distinct compositional changes. Sub-clustering identified diverse conventional and innate-like T cell subsets, including NKT, γδ T, and mucosal-associated invariant T (MAIT) cells (Fig. 2b; Supplementary Table 2). Of these, only MAIT cells were significantly increased by EAA treatment (Fig. 2c). MAIT and γδ17 cells occupied neighboring regions in UMAP space and were transcriptionally similar, as previously reported^20,40,41^. However, MAIT cells were highly enriched for αβ T cell-associated genes (e.g., *Trac*, *Lck*) and type 1/cytotoxic markers (e.g., *Ifng*, *Nkg7*), distinguishing them from γδ17 cells, which expressed *Trdc*, *Tcrg-V6*, *Cd163l1*, *Cxcr4*, and *Il17a* (Extended Data Fig. 2b,c).

**Figure 2.**
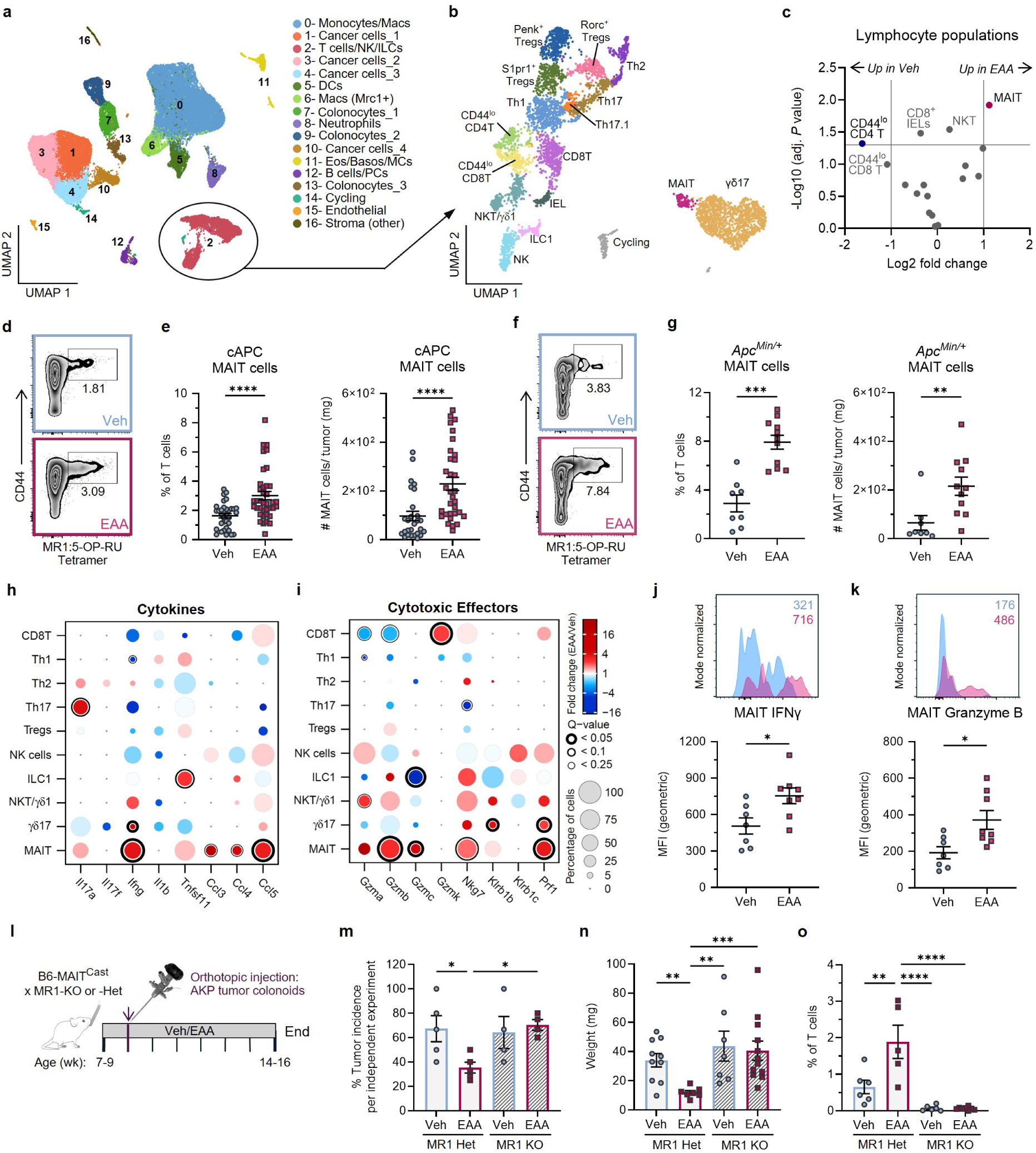
EAA increases cytotoxic tumor MAIT cells and anti-tumor effects are MAIT-dependent. **a**, UMAP plot of cells from cAPC tumors, colored by inferred cell types. Cluster 2, containing T cells, NK cells, and ILCs, is circled. **b**, UMAP of T cells, NK cells, and ILCs, labeled by inferred cell type. **c**, Volcano plot of populations from b, showing log_2_ fold change (log_2_FC) and -log_10_ adjusted *P* values following treatment. Populations are labeled if log_2_FC > 1 or adjusted *P* < 00.05. **d**, Representative flow cytometry plots of T cells from cAPC tumors gated on MR1 tetramer^+^ MAIT cells, with proportion of T cells indicated. **e**, Proportion (left) and number (right) of MAIT cells in cAPC tumors (n = 34-38 / group). **f**, Representative flow cytometry plots of T cells from *Apc*^Min/+^ tumors gated on MR1 tetramer^+^ MAIT cells. **g**, Proportion (left) and number (right) of MAIT cells in *Apc*^Min/+^ tumors (n = 8-11 / group). Dot plots showing differential expression of transcripts for cytokines (**h**) and cytotoxic molecules (**i**) by effector lymphocyte populations. Dot size indicates proportion of expressing cells, color indicates log_2_FC, and outline width reflects adjusted *P* (*Q*) value. **j**, Representative histogram and quantification of IFNγ expression in cAPC tumor MAIT cells (n = 7-8 / group). **k**, Representative flow cytometry histogram of granzyme B expression in cAPC tumor MAIT cells, with Mean Fluorescence Intensity (MFI) shown (top) and granzyme B MFI quantified (bottom). **l**, Experimental design for orthotopic injection of AKP tumor colonoids in B6-MAIT^Cast^ × MR1-KO or MR1-Het mice; vertical lines indicate weeks. **m**, Tumor incidence in five independent orthotopic AKP tumor experiments with MR1-KO/Het mice. Each data point represents tumor incidence from an independent experiment. **n**, Tumor weight of orthotopic AKP tumors (n = 5-8 / group). Each symbol represents data from an individual mouse. **o**, Proportion of AKP tumor T cells that were MR1:5- OP-RU tetramer^+^ MAIT cells (n = 5-8 / group). Data from six independent experiments (**d,e**), three (**f**,**g**), two (**j,k**), and five (**m**-**o**). Lines and error bars represent mean ± s.e.m. Statistical significance was determined by Wilcoxon rank-sum test with Benjamini-Hochberg FDR procedure (**c**,**h**,**i**), Mann-Whitney *U* test (**e**,**g**,**j**,**k**), or Kruskal-Wallis test (**n**,**o**). \**P* < 0.05, \*\**P* < 0.01, \*\*\**P* < 0.001, \*\*\*\**P* < 0.0001. Abbreviations: Basos (basophils), CD4T (CD4^+^ T cells), CD8T (CD8^+^ T cells), DCs (dendritic cells), Eos (eosinophils), IEL (intra-epithelial lymphocytes), IFNγ (interferon gamma), ILC (innate lymphoid cell), Macs (macrophages), MAIT (mucosal associated invariant T cell), MCs (mast cells), MFI (mean fluorescent intensity), NK (natural killer cells), NKT (natural killer T cells), PCs (plasma cells), Th (T helper), Tregs (regulatory T cells), UMAP (uniform manifold approximation and projection), and γδ1/17 (gamma delta 1/17 cells).

We validated the scRNA-seq findings by flow cytometry using MR1:5-OP-RU tetramers (gating in Extended Data Fig. 3; antibodies in Supplementary Table 3). Cytometry confirmed that EAA increased the proportion and number of MAIT cells in tumors of cAPC and *Apc^Min/+^* mice (Fig. 2d-g). Aligned with the transcriptomic data, this increase was specific to MAIT cells and not observed in other T cell populations (Extended Data Fig. 2d). The effect of EAA on extra-tumoral MAIT cells was more muted, with fewer MAIT cells detected in the adjacent colon or mesenteric lymph nodes (mLN; Extended Data Fig. 2e).

To compare EAA immune regulation with βHB, we administered EβHB and found no effect on tumor-infiltrating MAIT cells, despite an increase in serum βHB levels (Extended Data Fig. 1n,o; Extended Data Fig. 2f). EβHB altered other intra-tumoral T cell populations however, decreasing Th17 and increasing CD8 T cells (Extended Data Fig. 2f). These βHB effects on conventional T cells are consistent with prior studies^7,13,42^, but did not correspond with reduced tumor burden in cAPC mice (Extended Data Fig. 1p,q).

Having observed increased intra-tumoral MAIT cells, we next assessed whether EAA altered their effector functions. In EAA-treated tumors, MAIT cells upregulated *Ifng*, *Ccl5*, *Gzmb*, and *Gzmc*, suggesting enhanced cytotoxic potential (Extended Data Fig. 2c). Analysis across lymphocyte populations (annotated in Fig. 2b) showed that MAIT cells had the most significant increase in *Ifng*, *Gzmb*, *Gzmc*, *Nkg7*, and *Prf1* in EAA-treated tumors (Fig. 2h,i). Flow cytometry confirmed increased IFNγ and granzyme B protein levels in tumor-infiltrating MAIT cells, with a trend toward higher TNFα (Fig. 2j,k; Extended Data Fig. 2g,h,j). These effects were not observed in γδ, CD4, or CD8 T cells (Extended Data Fig. 2i,k). Thus, EAA selectively expands tumor-infiltrating MAIT cells and enhances their anti-tumor potential.

### MR1 is required for the anti-tumor MAIT cell effects of EAA

To test if MAIT cells are required for the anti-tumor effects of EAA, we used B6-MAIT^Cast^ mice, which possess an expanded MAIT cell compartment relative to C57BL/6 mice^33,43^. MR1 sufficient (MR1-Het) or MR1 deficient (MR1-KO) B6-MAIT^Cast^ littermates were treated with EAA or vehicle control and subjected to orthotopic transplantation of AKP tumor colonoids via colonoscopy (Fig. 2l)^31,39,44^. EAA reduced tumor incidence in MR1-sufficient mice whereas tumor incidence was not reduced in MR1-deficient mice regardless of treatment (Fig. 2m). Similarly, EAA reduced tumor mass relative to vehicle in MR1-sufficient mice, with no effect observed in MR1-deficient mice (Fig. 2n). Consistent with our observations in the GEMM CRC models, EAA increased tumor-infiltrating MAIT cells in MR1-sufficient mice, whereas MR1-deficient tumors contained few MAIT cells irrespective of treatment (Fig. 2o). These data establish that MR1 and MAIT cells are essential for the anti-tumor effects of EAA and highlight the MR1-MAIT axis as a tractable therapeutic target.

### AcAc and 5-A-RU expand human MAIT cells and enhance their cytotoxicity in an MR1-dependent manner

The enhancement of anti-tumor MAIT cell responses by exogenous AcAc raises the question of how treatment mediates this effect. To address this, we cultured human peripheral blood mononuclear cells (PBMCs) with AcAc or controls and assessed MAIT cell expansion (Fig. 3a; gating in Extended Data Fig. 5; antibodies in Supplementary Table 4)^45^. Neither AcAc, βHB, nor the reactive dicarbonyl MGO alone promoted MAIT cell expansion relative to NaCl (Fig. 3b). The riboflavin intermediate 5-A-RU and MGO form the MAIT activating MR1 ligand 5-OP-RU^25–27^; so as expected, supplementation of cultures with 5-A-RU and MGO induced MAIT cell expansion. Notably, co-administration of 5-A-RU and AcAc induced dose-dependent MAIT cell expansion (Fig. 3b), consistent with our in vivo findings. Addition of anti-MR1 blocking antibodies abrogated MAIT cell expansion in both the MGO and AcAc plus 5-A-RU conditions (Fig. 3c). We used PBMCs from a second donor to confirm that AcAc mediated MR1-dependent MAIT expansion was not a donor-dependent epiphenomenon (Extended Data Fig. 4a-c; gating in Extended Data Fig. 5).

**Figure 3.**
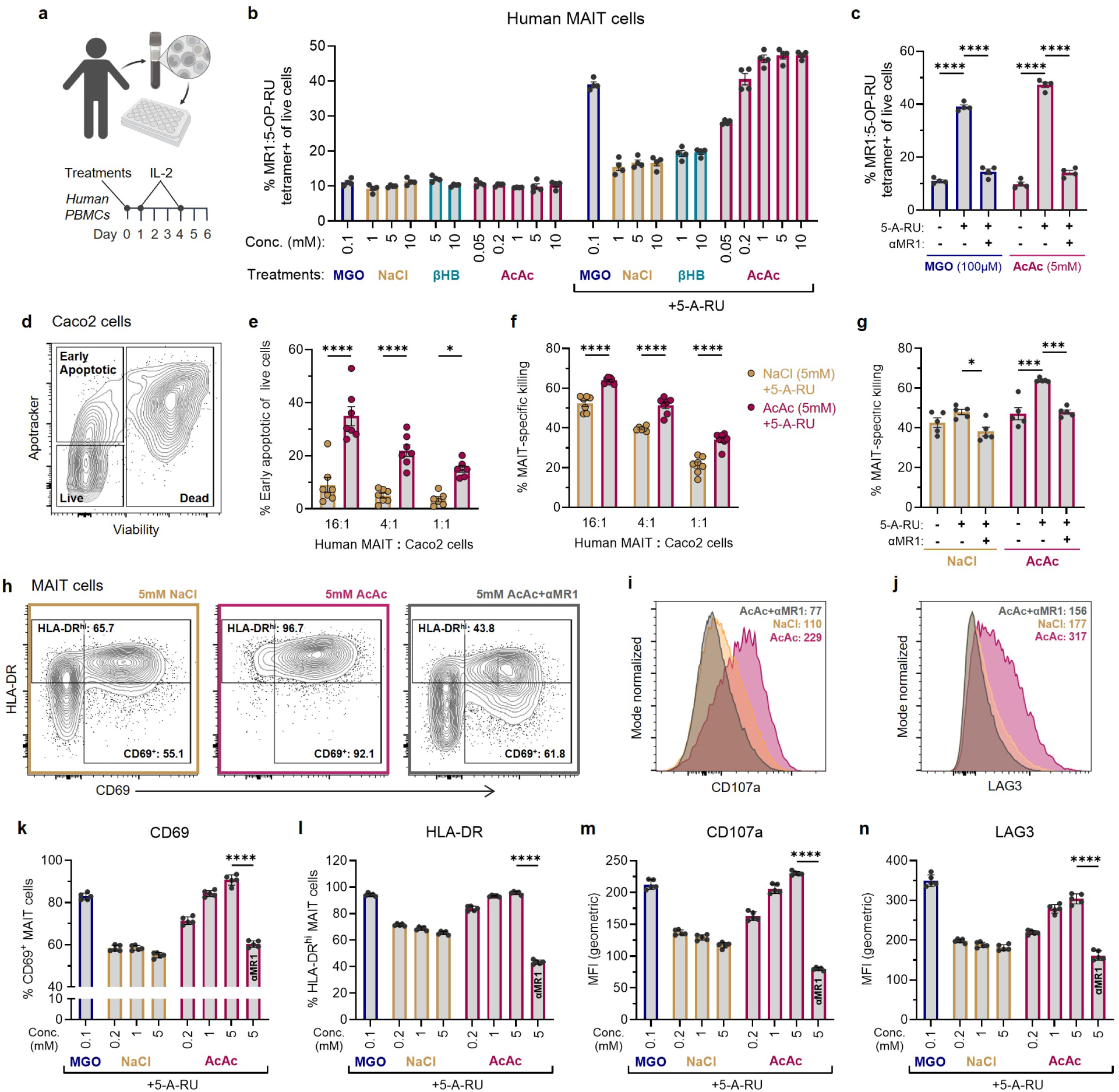
AcAc expands human MAIT cells and enhances tumor cell killing. **a**, Schematic representation of *in vitro* expansion of human MAIT cells from peripheral blood mononuclear cells (PBMCs). **b**, Proportion of PBMCs (donor 24A) that were MR1 tetramer^+^ MAIT cells after 6 d of culture with the indicated treatments (n = 4 / group). **c**, Proportion of MR1 tetramer^+^ MAIT cells following culture with methylglyoxal (MGO) or AcAc, with or without 5-A-RU and αMR1 antibody (n = 4 / group). **d**, Representative flow cytometry plot from human MAIT-Caco2 co-cultures showing Caco2 cells that are live, early apoptotic, or dead. **e**, Proportion of live Caco2 cells that were Apotracker^+^ (early apoptotic) after 24 h co-culture with MAIT cells (n = 7 / group). **f**, Proportion of dead Caco2 cells following MAIT cell co-culture, adjusted by subtracting the proportion of cell death in matched Caco2-only cultures (n = 7 per group). **g**, Proportion of dead Caco2 cells after 24 h of culture with MAIT cells (4:1 MAIT:Caco2) and treatments with 5 mM NaCl, 5 mM AcAc, or 100 μM MGO, with or without 5-A-RU and αMR1 antibody; values adjusted as in (f) (n = 5 per group). **h**, Representative flow cytometry plots of CD69 and HLA-DR expression by human MAIT cells cultured for 24 h with Caco2 cells (4:1 ratio) and indicated treatments; proportions shown represent total cells. Representative histograms of CD107a (**i**) and LAG3 (**j**) expression by MAIT cells under the same culture conditions; geometric mean fluorescence intensity (MFI) is indicated. **k**, Proportion of MAIT cells that were CD69^+^. **l**, Proportion of MAIT cells that were HLA-DR^hi^. **m**, Geometric MFI of CD107a by MAIT cells. **n**, Geometric MFI of LAG3 by MAIT cells following 24 h of culture with Caco2 cells and indicated treatments; αMR1 conditions indicated within bars (**k**-**n**). Data are representative of three independent experiments (**b,c**) or two independent experiments (**e**,**f, g**-**n**). Lines and error bars represent mean ± s.e.m. Statistical significance was determined by one-way ANOVA with Tukey’s multiple comparison test (**c**,**g**), two-way ANOVA with Tukey’s multiple comparison test (**e**,**f**), or one-way ANOVA with Šídák’s multiple comparisons test (**k**-**n**). \**P* < 0.05, \*\*\**P* < 0.001, \*\*\*\**P* < 0.0001.

To assess MAIT-mediated tumor cell killing, expanded and bead-enriched human MAIT cells were co-cultured with Caco2 cells, a human colonic adenocarcinoma cell line (Extended Data Fig. 4d,e). After 24 hours, live, early apoptotic, and dead Caco2 cells were identified using flow cytometry (Fig. 3d). Co-cultures treated with AcAc plus 5-A-RU increased early apoptotic Caco2 cells and MAIT-specific killing across several MAIT:Caco2 ratios (Fig. 3d-f); these effects were abolished by anti-MR1 antibody (Fig. 3g).

To identify if increased tumor cell killing could be attributed to changes in MAIT cell activation or cytotoxicity, we measured CD69, HLA-DR, LAG3 and CD107a (LAMP-1) expression in MAIT cells by flow cytometry (Fig. 3h-n). MAIT-Caco2 co-cultures with 5-A-RU plus AcAc showed dose-dependent upregulation of each marker, which was blocked by anti-MR1 antibody. Collectively, these results demonstrate that AcAc and 5-A-RU drive MR1-dependent human MAIT cell expansion, activation, and tumor cell killing.

### AcAc and 5-A-RU upregulate surface MR1 expression in tumor-infiltrating monocytes, human PBMCs, and CRC cell lines

As AcAc-driven MAIT cell expansion, activation, and tumor killing are MR1-dependent, we hypothesized that AcAc enhances MR1 antigen presentation, which requires surface expression of MR1. Consistent with this, both MGO and AcAc with 5-A-RU rapidly increased surface MR1 expression on total human PBMCs, as assessed by mean fluorescence intensity (MFI) and the proportion of MR1^hi^ cells (Fig. 4a; Extended Data Fig. 6a,b). PBMCs comprise heterogeneous cell types (Extended Data Fig. 6c; Supplementary Table 5), prompting our analysis of specific populations to identify those upregulating MR1 in response to AcAc or MGO with 5-A-RU. Several cell types increased surface MR1 (Fig. 4b,c; Extended Data Fig. 6d-f), but with distinct kinetics. Most PBMCs, including dendritic cells (DCs), which are well-characterized antigen presenting cells, exhibited a modest and transient increase in MR1 that peaked within 24 h (Fig. 4b, Extended Data 6d-f). However, monocytes displayed a more intense and sustained response (Fig. 4c), suggesting they may promote human MAIT cell activation and expansion. Next, we assessed the effect of AcAc and 5-A-RU on MR1 expression by Caco2 cells. AcAc or MGO plus 5-A-RU increased surface MR1 expression, which peaked after 4-5 h, whereas MHC class I (HLA-A, B, C) expression remained unaffected (Extended Data Fig. 6g-i).

**Figure 4.**
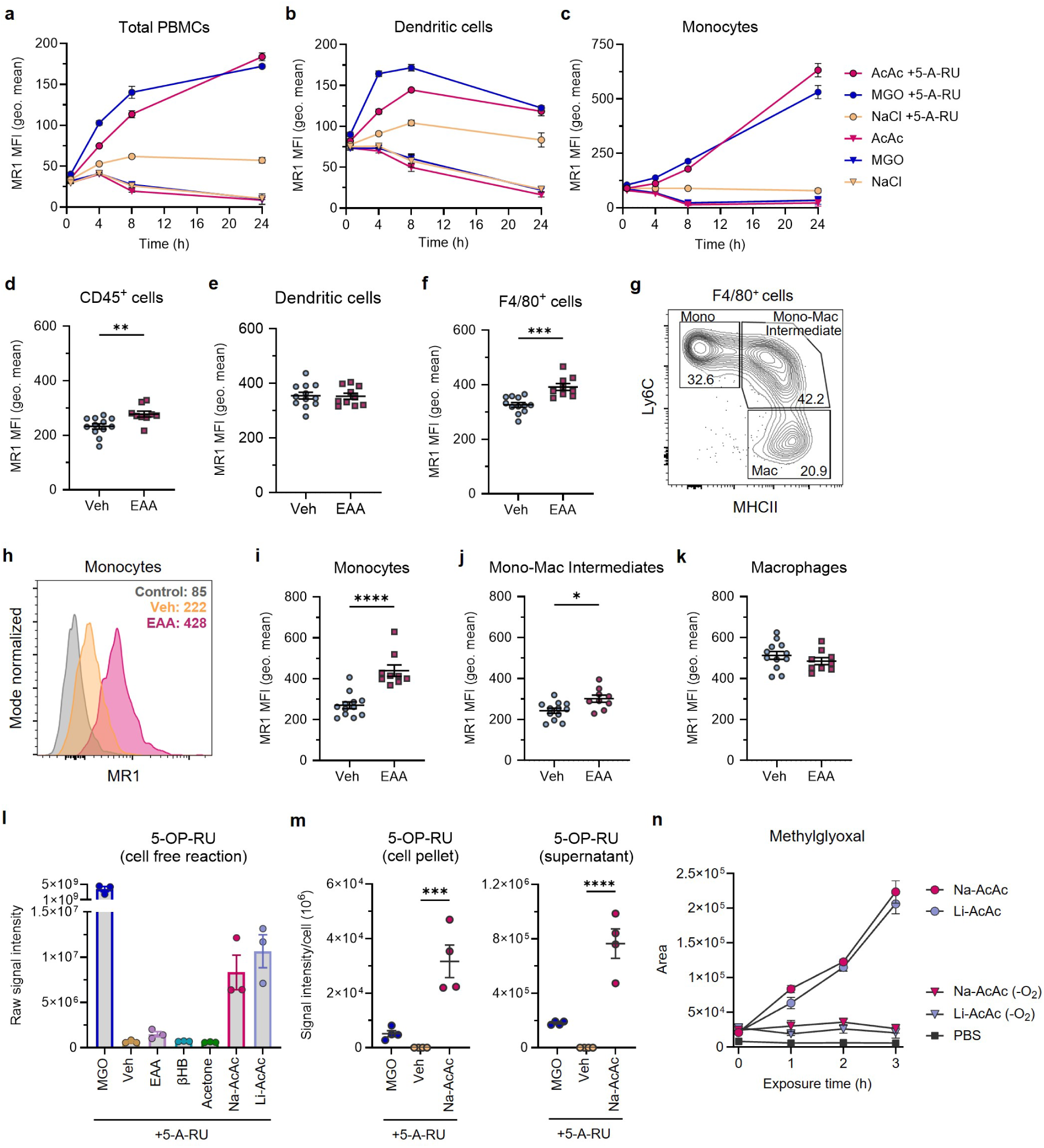
AcAc and 5-A-RU increase surface MR1 and generate 5-OP-RU. **a**-**c**, Time course of surface MR1 expression (geometric MFI) in total human PBMCs (**a**), dendritic cells (**b**), or monocytes (**c**), cultured with 5mM AcAc, 5mM NaCl, or 100μM methylglyoxal (MGO) with or without 5-A-RU (n = 5 / group). At 24 h, AcAc/MGO+5-A-RU vs AcAc/MGO: *P* < 0.0001 (**a**,**c**) and *P* < 0.01 (**b**). **d**-**f**, Surface MR1 expression of cAPC tumor CD45+ cells (**d**), dendritic cells (**e**) or F4/80+ cells (**f**) (n = 9-12 / group). **g**, Representative flow cytometry plot of F4/80+ cells gated as Ly6C^hi^ MHC II^−^ monocytes, Ly6C^hi^ MHC II^+^ monocyte-macrophage intermediates, or Ly6C^low^ MHC II^+^ macrophages. **h**, Representative histograms of surface MR1 (geometric MFI) by P1 monocytes from cAPC tumors. **i**-**k**, Surface MR1 staining of cAPC tumor F4/80+ subsets (n = 9-12 / group). **l**, Measurement of 5-OP-RU by liquid chromatography-mass spectrometry (LC-MS) following 30 min cell free reactions with 5-A-RU plus MGO, vehicle (Veh), ethyl acetoacetate (EAA), β-hydroxybutyrate, acetone, Na-AcAc, or Li-AcAc (n = 3 / group). **m**, 5-OP-RU measurement from cell pellets (left) or supernatants after 3 h PBMC cultures with 5-A-RU plus MGO, Veh, or Na-AcAc (n = 4 / group). **n**, LC-MS detection of MGO generated from Li-AcAc, Na-AcAc or PBS exposed to atmospheric oxygen or incubated under anaerobic (-O_2_) conditions (n = 3 / group). At 3 h, Na/Li-AcAc vs Na/Li-AcAc (-O_2_): *P* < 0.01. Data are representative of two independent experiments (**a**-**c,m**) or three independent experiments (**d**-**f**,**i**-**l,n**). Lines and error bars represent mean ± s.e.m. Statistical significance was determined by two-way ANOVA with Geisser-Greenhouse correction and Tukey’s multiple comparison test (**a**-**c**), Mann-Whitney *U* test (**d**-**f**,**i**-**k**) or one-way ANOVA with Tukey’s multiple comparison test. \**P* < 0.05, \*\**P* < .01, \*\*\**P* < 0.001, \*\*\*\**P* < 0.0001.

To determine if exogenous AcAc regulates surface MR1 expression in vivo, we analyzed MR1 levels in cAPC tumors. Due to technical challenges of staining murine MR1 with monoclonal antibodies, we processed and stained tumor cells with a biotinylated polyclonal anti-MR1 antibody. EAA treatment increased MR1 expression in tumor-infiltrating immune cells, but not in EpCAM^+^ tumor cells (Fig. 4d; Extended Data Fig. 6j). Further analysis of immune cells revealed no change in MR1 expression on DCs (Fig. 4e), but a marked increase in F4/80^+^ cells (Fig. 4f), comprising Ly6C^hi^ MHCII^−^ monocytes, Ly6C^hi^ MHCII^+^ monocyte-macrophage intermediates, and Ly6C^low^ MHCII^+^ macrophages (Fig. 4g)^46,47^. Of these, the largest increase in MR1 expression occurred in Ly6C^hi^ MHCII^−^ monocytes, with no significant change in macrophages (Fig. 4h-k). MHCI levels remained unchanged across all populations analyzed (Extended Data Fig. 6k-s). In summary, surface MR1 expression was rapidly upregulated by AcAc plus 5-A-RU in human cells and by EAA in tumor-infiltrating immune cells in mice. In both contexts, monocytes showed the most pronounced increase in MR1.

### AcAc and 5-A-RU generate a potent MAIT cell-activating MR1 ligand

Previous studies have shown that surface MR1 expression is regulated by ligand availability^25,48,49^, prompting the hypothesis that AcAc and 5-A-RU generate an MR1 ligand. We tested whether AcAc and 5-A-RU generate the MAIT activating MR1 ligand 5-OP-RU, a pathway has not been previously described. Remarkably, both sodium acetoacetate (Na-AcAc, prepared by hydrolysis of EAA and used in Fig. 3 of this study) and high-purity lithium acetoacetate (Li-AcAc) reacted spontaneously with 5-A-RU to generate 5-OP-RU in cell-free reactions (Fig. 4l). In time course experiments, we found that AcAc and 5-A-RU are rapidly consumed as 5-OP-RU is generated, and 5-OP-RU is absent in reactions lacking AcAc (Extended Data Fig. 7a,b). In addition to 5-OP-RU, its degradation product 7-methyl-8-D-ribityllumazine (RL-7-Me) was detected in MGO or AcAc plus 5-A-RU conditions (Extended Data Fig. 7c).

Due to their instability, metabolites such as MGO, 5-A-RU, and 5-OP-RU are difficult to detect in tissue samples such as tumors^25,27^. To determine if 5-OP-RU could be generated in a cell-based system, we incubated human PBMCs with AcAc, NaCl, or MGO in the presence of 5-A-RU. Following a 3 h culture with AcAc plus 5-A-RU, 5-OP-RU was detected in both cell pellets and culture supernatants (Fig. 4m). RL-7-was also present in supernatants (Extended Data Fig. 7d), and AcAc was readily detected in both supernatants and cell pellets (Extended Data Fig. 7e). These findings suggest that 5-OP-RU is either taken up by cells from the culture medium or formed intracellularly from AcAc and 5-A-RU.

The unexpected formation of 5-OP-RU from AcAc and 5-A-RU led us to hypothesize that AcAc generates methylglyoxal (MGO), the reactive carbonyl that converts 5-A-RU to 5-OP-RU. Because this reaction occurred in cell-free conditions, we proposed that it proceeds non-enzymatically in the presence of oxygen. To test this hypothesis, we performed cell-free reactions incubating AcAc in the presence of atmospheric oxygen or under anaerobic conditions, and measured MGO formation over time (Fig. 4n). Spontaneous, non-enzymatic generation of MGO from AcAc was observed only under ambient oxygen conditions, as detected by derivatization with 4-methoxy-o-phenylenediamine (4-PDA) (Extended Data Fig. 7f-k). Together, these results support that AcAc generates MGO and subsequently 5-OP-RU when 5-A-RU is available (Extended Data Fig. 7l).

## Discussion

In this study, we demonstrate that oral administration of EAA elevates serum and tumor AcAc, suppressing tumor development and progression in multiple colorectal cancer models via MAIT cells. Due to the instability of AcAc, previous studies have primarily used βHB^5,12,13^; however, our results show that EAA but not EβHB suppressed tumor burden, highlighting the importance of investigating ketone-specific effects. AcAc’s inherent reactivity promoted anti-tumor effects via a previously unreported non-enzymatic generation of MGO, which results in 5-OP-RU in the presence of 5-A-RU, fueling the expansion and activation of cytotoxic MAIT cells.

Prior reports have described both pro- and anti-tumorigenic roles for MAIT cells^28–32^. Our data show that exogenous AcAc expands and activates cytotoxic MAIT cells to promote tumor control in GEMM and orthotopic CRC models. While immune, epithelial, and tumor cells can present antigen via MR1 to activate MAIT cells^21,24,32^, our results identify monocytes as important MR1 antigen presenting cells following AcAc treatment of human PBMCs and in EAA-treated cAPC tumors.

Our observations that AcAc generates MGO in the presence of oxygen, and 5-OP-RU subsequently forms when microbially derived 5-A-RU is available, is a plausible scenario within the colonic tumor microenvironment (TME). Colonic tumors are vascularized, albeit with lower oxygen tension than healthy tissues, and the TME can be enriched for reactive oxygen species^50,51^, suggesting multiple potential catalysts to convert AcAc to MGO. The hyperpermeable vasculature of tumors as opposed to normal colonic tissues^52,53^, may contribute to the accumulation of AcAc in tumors vs normal tissue (Fig 1c), and subsequent enhanced MR1 antigen generation underlying the increased MAIT cell expansion observed in tumor versus normal tissue (Fig 2e,g,o). Intra-tumoral MGO could proceed to generate 5-OP-RU since colonic tumors are exposed to microbial metabolites from luminal and intra-tumoral bacteria, including MR1 ligand precursors^34,54^. Collectively, these TME features may favor the generation of MGO from exogenous AcAc to drive 5-OP-RU generation and boost MAIT cell driven anti-tumor immunity.

Limitations of this study include the technical challenge of directly quantifying highly reactive and unstable metabolites such as 5-A-RU or 5-OP-RU in tissues. Assessment of 5-OP-RU in tumors relied on indirect measures such as MR1 expression and MAIT cell expansion; however, successful direct detection in cell cultures treated with AcAc and 5-A-RU demonstrates intracellular uptake or generation of 5-OP-RU. Together, these findings support that 5-OP-RU is generated from AcAc and 5-A-RU in biological systems.

Future studies are needed to determine if exogenous AcAc offers clinical benefit for CRC patients or for individuals with colonic polyps and at high risk for CRC, and whether the MR1-MAIT axis can be targeted in human tumors. The potential for AcAc to be used therapeutically in other cancers, as well as in combination with immunotherapies such as checkpoint inhibitors, also warrants further investigation. Finally, the effects of exogenous AcAc on tumor and immune metabolism, independent of MR1 ligand generation, remain to be determined. Delineating any metabolic effects could improve our mechanistic understanding of the safety and efficacy of exogenous AcAc.

In summary, these results establish exogenous AcAc as an accessible intervention that promotes anti-tumor MAIT cells by generating the canonical MR1 ligand 5-OP-RU and identify the MR1-MAIT axis as a potential target for CRC immunotherapy.

## METHODS

### Mice

C57BL/6, *Cdx2*^Cre^ x *Apc*^flox/+^ (cAPC), *Apc^Min/+^,* B6*-*MAIT^Cast^, and B6*-*MAIT^Cast^ x *Mr1*^−/−^ mice were bred and housed at the Harvard T.H. Chan School of Public Health (HSPH) barrier facility under specific pathogen free conditions, with 12 h day/night cycles. C57BL/6J (JAX 0000664), *Cdx2*^Cre^ (JAX 009350), *Apc*^flox/+^ (JAX 009005), and *Apc^Min/+^* (JAX 002020) mice were initially purchased from Jackson Laboratory. B6*-*MAIT^Cast^, and B6*-*MAIT^Cast^ x *Mr1*^−/−^ mice were provided by Prof. Mansour Haeryfar (University of Western Ontario, CA). Male and female mice were used, and mice were assigned to treatment groups to ensure even distribution of sex, age, and litter of origin across experimental conditions. Animal studies were approved and conducted in accordance with Harvard Medical School’s Standing Committee on Animals and NIH guidelines.

### In vivo treatments

#### Administration of ketone esters

Ethyl acetoacetate (EAA; ThermoFisher Scientific, 20 mM) or vehicle only control was administered in the drinking water and provided ad libitum. EAA was administered to cAPC mice from 6-8 wk of age, except in experiments in mice with established tumors, where EAA was provided from 14-16 wk. For *Apc^Min/+^* mice, EAA was started 1 wk prior to dextran sulfate sodium (DSS) treatment, or 1 wk before colonoscopy-guided tumoroid injection in B6-MAIT^cast^ mice (see below). For β-hydroxybutyrate ester (EβHB) studies, d-β-hydroxybutyrate-(R)-1,3-butanediol monoester (KetoneAid) or vehicle control was added to drinking water at 20 mM from 6-8 wk of age until experimental endpoint. All drinking water treatments were refreshed twice weekly. Vehicle controls were water for EAA and water containing matched quantities of stevia extract, citric acid, allulose, and potassium sorbate (Stevia Liquid Drops, Natrisweet) for EβHB.

#### Administration of dextran sulfate sodium (DSS) to Apc^Min/+^ mice

3% DSS (ThermoFisher Scientific) was added to the drinking water of *Apc^Min^*^/+^ mice to promote development of colonic adenocarcinomas^38^. DSS-treated water was provided ad libitum for four days, four weeks prior to tissue collection.

#### Colonoscopy-guided orthotopic tumor transplantation

For orthotopic tumor transplantations, tumor colonoids (cultured as described below) were dissociated with TrypLE Express (Thermo Fisher Scientific) for 5 min at 37°C, resuspended in basal medium containing 10% Cultrex Reduced Growth Factor Basement Membrane Matrix (BME; R&D Systems) and injected into the colonic mucosa of recipient mice (8-12 weeks old) by colonoscopy as previously described^39^. Briefly, a Karl Storz endoscope system consisting of a documentation system (AIDA), cold light source (Xenon 175), imaging system (Image1 S; all Karl Storz), and a Hopkins telescope (1232 AA) were used. Mice were anesthetized with isoflurane, and colonoids were injected using a Hamilton syringe, transfer needle, and an injection custom needle (Hamilton Inc.). The injection needle was placed on the colonic mucosa with the bevel facing the tissue, and 100 µL of medium containing 150,000 organoid cells was injected to form a mucosal bubble. Tumors were harvested 4-6 weeks post-injection.

### Ketone measurement in murine samples

#### Sample extraction and derivatization

Fecal pellets or tissues (25-60 mg) were homogenized in bead beater vials (Qiagen; fecal samples with 2.4 mm steel beads; tissue with a garnet matrix) and mixed with 300 µl internal standard (IS; 2 µM 3-hydroxybutyrate-d₄ or acetoacetate-d₄ in acetonitrile) and 300 µl water using a TissueLyser II (Qiagen) for 10 min at 50 Hz. Serum (100 µl) was mixed with water to a total of 300 µl and mixed with 300 µl IS. All samples were centrifuged at maximum speed (10 min, 4°C) and 560 µl supernatant was collected. Extracts were derivatized by adding 40 µl 2-nitrophenylhydrazine (200 mM in 50% acetonitrile in water) and 40 µl EDC (120 mM in 50% acetonitrile and 6% pyridine in water), then incubating 30 min at 40°C. After centrifugation, supernatants were transferred to LC-MS vials. Standard curves were prepared from ketone standards (100 µM each, 1:5 serial dilutions in water) and processed identically to samples.

#### Liquid chromatography-mass spectrometry (LC-MS) analysis

LC-MS was performed as previously described^55^. Derivatized samples were analyzed on a ThermoFisher Q Exactive Plus Orbitrap mass spectrometer (HESI source, Ultimate 3000 LC). Separation was by Kinetex C18 EVO (2.6 µm, 150 × 2.1 mm) at 30°C (4°C autosampler), injecting 1-2 µl. The mobile phase was (A) water with 0.1% formic acid and (B) acetonitrile with 0.1% formic acid, using the following gradient: 5% B (0-5 min), to 100% B (25-30 min), hold at 100% B (30.0-30.01 min), re-equilibrate to 5% B (35 min); flow 0.2 mL min⁻¹ (0-30 min), 0.37 mL min⁻¹ (30-35 min). LC flow was diverted to waste at 0-8.8 and 30-35 min. MS full scan settings: 70,000 resolution, 1 × 10⁶ AGC target, 50 ms max IT, 160-600 m/z.

### Glucose measurement

Blood glucose was measured from whole blood collected by cardiac puncture. Mice were euthanized and blood withdrawn using a 26G needle. 2-5 µl of freshly collected blood was applied directly to a Precision Xtra glucose test strip (Abbott), and glucose concentration was recorded using the corresponding Precision Xtra meter, following manufacturer’s instructions.

### Histopathological assessment

Colons were opened longitudinally and luminal contents removed. Tissues were fixed in 4% paraformaldehyde, transferred to 70% ethanol, and subsequently paraffin-embedded and sectioned by the Dana-Farber/Harvard Cancer Center Rodent Histopathology Core (Boston, MA). Five hematoxylin and eosin (H&E)-stained sections per sample were assessed by a board-certified gastrointestinal pathologist (J.N.G.) who was blinded to the experimental conditions and groups. Slides were examined for the presence of aberrant crypt foci, adenoma, or adenocarcinoma.

### Tissue processing

#### Cell isolation from colonic tumors

Colonic tumors were excised, washed in cold phosphate-buffered saline (PBS), weighed, and chopped into pieces <1 mm in RPMI 1640 (ThermoFisher Scientific) supplemented with 3% fetal bovine serum (FBS; Corning) and 1% penicillin/streptomycin (Corning). Tissue fragments were transferred to 10 mL digestion buffer [RPMI 1640 with 3% FBS, 0.5 mg/mL collagenase D (Roche), 0.03 mg/mL DNase I (Roche), 0.5 mg/mL dispase (STEMCELL Technologies), and 1% penicillin/streptomycin], and incubated at 37 °C for 45 min on a rotating mixer. Digests were vortexed and filtered through 70 µm cell strainers (VWR) into PBS containing 5 mM EDTA and 2% FBS. Cells were washed, resuspended in FACS buffer (PBS with 2% FBS and 1 mM EDTA), and counted using a LUNA-FX7 cell counter (Logos Biosystems) prior to downstream applications.

#### Cell isolation from colonic lamina propria

Colons were excised, trimmed of visible fat and vasculature, opened longitudinally, and washed in PBS. Tissues were incubated for 10 min on ice in PBS with 5 mM HEPES and 1 mM dithiothreitol (DTT; both ThermoFisher Scientific). Samples were then incubated twice for 15 min at 37°C on a rotating mixer in PBS containing 5 mM EDTA (Millipore Sigma) and 3% FBS (ThermoFisher Scientific). Tissues were then rinsed in PBS, chopped into pieces <1 mm, and digested in the enzyme cocktail described for tumor samples, using two 20 min cycles at 37°C on a rotating mixer. Cell suspensions were strained, washed and counted as described above.

#### Cell isolation from mesenteric lymph nodes (mLN)

mLNs were excised and cleared of fat and connective tissue, then chopped into pieces <1 mm in RPMI 1640 containing 3% FBS and 1% penicillin/streptomycin (Corning). Suspensions were transferred to 4 mL digestion buffer [RPMI 1640 with 3% FBS, 0.25 mg/mL Liberase TL (Sigma-Aldrich), 0.5 mg/mL collagenase D, 0.03 mg/mL DNase I, and 1% penicillin/streptomycin], and digested for 25 min at 37°C on a rotating mixer. Digests were vortexed and filtered through 70 µm strainers into PBS containing 5 mM EDTA and 2% FBS. Single-cell suspensions were washed and counted as described above.

### scRNA-Seq

scRNA-seq samples were prepared using the Chromium Next GEM Single Cell Fixed RNA Kit (10x Genomics), which enables sample fixation, freezing, and pooling. Tumor samples from five independent experiments involving 40 mice were analyzed, yielding 35 tumors from 16 tumor-bearing mice.

#### Sample preparation

Colon tumors were excised and processed into single-cell suspensions as described above. Ly6G^+^ neutrophils are the most abundant population in cAPC tumors, so samples were de-enriched for neutrophils to allow greater sequencing read depth of other cell types. After enzymatic digestion, 90% of each sample was labelled with Anti-Ly6G UltraPure Microbeads (Miltenyi Biotec) and Ly6G^+^ cells were depleted according to manufacturer’s instructions. Neutrophil-depleted samples were then added back to their corresponding untreated fraction. Pre-fixation sample cell counts (2.5 × 10^5^ −5.4 × 10^6^) and viability (80% - 90%) were determined using a LUNA-FX7 cell counter. Samples were fixed and stored at −80 °C, using Chromium Next GEM Single Cell Fixed RNA Sample preparation kit (10x Genomics), according to manufacturer’s instructions.

#### Library construction and sequencing

To avoid batch effects, all 16 samples were thawed, normalized by cell count, and allocated into groups, according to treatment and sex (3.0 × 10^5^ −1.0 × 10^6^ cells / group), and each group was barcoded for multiplexing at the same time. Probe hybridization of pooled samples was performed in two independent reactions with Chromium Fixed RNA Kits: Mouse Transcriptome, 4 rxns x 4 BC (10x Genomics) according to manufacturer’s instructions. Gel Bead-in-Emulsion (GEMs) were generated from each reaction on a Chromium X instrument (10x Genomics) with a target of 40,000 cells recovered (10,000 / barcode), and libraries were prepared according to manufacturer’s instructions. Sequencing was performed on a NovaSeq S4 Sequencer (Illumina) with a target of 50,000 reads per cell. Library construction and sequencing was carried out at the Bauer Sequencing Core (Cambridge, MA).

#### scRNA-Seq data processing

Raw sequencing data were pre-processed using Cell Ranger Multi (v7.0.0; 10x Genomics) and aligned with the mouse mm10 reference transcriptome (Supplementary Table 6). Demultiplexing, barcode whitelisting, and read assignment were performed according to the Cell Ranger pipeline; only reads with valid sample indexes, cell barcodes, and unique molecular identifiers (UMIs) > 500 were retained for downstream analysis.

Subsequent processing, quality control, analysis, and visualization were performed using Cellenics (Harvard Medical School-hosted instance). A cell size filter was applied to exclude debris or cells with low UMI counts. The threshold was set at the inflection point of #UMI vs cell rank ‘knee’ plots. A mitochondrial content filter (3 median absolute deviations above median) and a linear threshold on gene vs UMI count regression (*P* < 0.0003) were applied to remove outliers. Doublets were identified and excluded using scDblFinder (Bioconductor). After filtering, a total of 37,402 cells were retained across conditions (median = 3,506 genes per cell).

After processing, gene expression data were log-normalized and integrated across samples using Harmony. Highly variable genes (n = 2,000) were identified by the ‘vst’ method. Dimensionality was reduced using principal component analysis (PCA, 28 components), and uniform manifold approximation and projection (UMAP) embedding (minimum distance = 0.3) and Louvain clustering (resolution = 0.8) were applied. Clusters were annotated with a semi-supervised approach, using lineage markers (e.g., T cells, B cells, myeloid cells, epithelial cells) and by comparing top differentially expressed genes (DEGs) in each cluster to published and publicly available datasets. Automated annotation tools (e.g., ScType-intestine-mouse and ScType-immune-system-mouse) ^56^ were used to cross-reference and validate annotation. DEGs were calculated using the Wilcoxon rank-sum test, and *P*-values were adjusted using the Benjamini-Hochberg procedure to control the false discovery rate (FDR). Top DEGs for each cluster are in Supplementary Table 1-2. Dot plots (Fig. 2h,i) were generated in R (v4.1.3) using ggplot2 (v3.5.1), data.table (v1.14.8), plyr (v1.8.9), and scales (v1.3.0) packages.

### Flow cytometry

#### Surface staining

Single-cell suspensions were incubated with LIVE/DEAD Fixable Near-IR Stain (ThermoFisher Scientific) and TruStain FcX (BioLegend) in PBS for 15 min on ice. Cells were then stained with antibodies in FACS buffer (PBS with 2% FBS and 1 mM EDTA) for 20 min on ice. Antibodies for murine and human samples are listed in Supplementary Table 3 and 4, respectively. For experiments using MR1 or CD1d tetramers (NIH Tetramer Core Facility), tetramers were added with surface antibodies and incubated in FACS buffer for 30 min at room temperature (rt). Cells were washed twice with FACS buffer and analyzed on a BD FACSymphony A3 Cell Analyzer (BD Biosciences) or fixed for intracellular staining.

#### Intracellular staining

For detection of intracellular markers, cells were fixed and permeabilized using the eBioscience Foxp3/Transcription Factor Staining Buffer Set (ThermoFisher Scientific) per manufacturer’s instructions. Samples were stored overnight at 4°C in permeabilization buffer, then stained with antibodies for 1 h at rt. After staining, cells were washed, resuspended in FACS buffer, and acquired on a BD FACSymphony A3 Cell Analyzer. For cytokine detection, cells were cultured in RPMI 1640 with 10% FBS and Protein Transport Inhibitor Cocktail (ThermoFisher Scientific) for 4 h at 37°C then fixed in BD Cytofix Fixation Buffer (BD Biosciences) for 30 min at rt. If nuclear targets were labeled, samples were then fixed and permeabilized using the eBioscience Foxp3/Transcription Factor Staining Buffer Set, followed by intracellular staining as above.

#### MR1 staining of mouse cells

For MR1 staining of murine samples, tissues were collected in FACS buffer containing rabbit polyclonal anti-MR1 antibody (FabGennix, #MR1-BIOTIN; 1:400), chopped and digested in enzyme buffer as described above, supplemented with anti-MR1 antibody (1:800). Surface staining was performed at 4°C for 30 min with anti-MR1 antibody (1:100). Following surface staining, samples were stained with APC-conjugated streptavidin (SA) in FACS buffer (0.5μg/mL) for 30 min on ice, to label surface MR1. Cells were fixed with BD Cytofix Fixation Buffer and permeabilized with the Foxp3/Transcription Factor Staining Buffer Kit. Samples were then incubated with BV421-conjugated SA in permeabilization buffer for 30 min on ice to label MR1 that was surface-localized during initial staining but internalized during processing. After SA labeling, intracellular antibody staining and analysis was performed as described above. Surface MR1 expression was quantified as the sum of geometric mean fluorescence intensities (MFI) of APC-SA and BV421-SA.

#### Flow cytometry analysis

Data was analyzed with FlowJo version 10.10 (BD Biosciences). Representative hierarchical gating strategies are in Extended Data Fig. 3 (mouse tumor cells) and Extended Data Fig. 5 (human cells).

### Acetoacetate synthesis

Acetoacetate (AcAc) for in vitro experiments was prepared by hydrolysis of EAA. 1mL EAA and 8mL 1M NaOH were mixed at 60 °C for 30 min, stirring. Samples were adjusted to pH 7.2 with 50% HCl and stored in the gaseous phase of liquid nitrogen. AcAc concentration was measured with the Autokit Total Ketone Bodies Assay (Fujifilm Wako).

### Processing human peripheral blood mononuclear cells (PBMCs)

Peripheral blood mononuclear cells (PBMCs) were isolated from fresh human Leukopaks (STEMCELL Technologies; Cat# 70500.2) within 12 h of collection and according to manufacturer’s instructions. Briefly, samples were diluted 1:1 with PBS containing 2% FBS, layered over Ficoll-Paque PLUS (GE Healthcare), and centrifuged at 800 × g for 20 min at rt, no brake. Cells at the interface were washed twice with PBS/2% FBS and centrifuged at 300 × g for 10 min. Platelets were removed in an additional wash and centrifugation at 120 x g for 10 min, brake off. Cell counts and viability were determined using a LUNA-FX7 cell counter prior to downstream applications. Purified PBMCs were used immediately or frozen for storage in FBS/20% dimethyl sulfoxide (DMSO) in liquid nitrogen. All Leukopaks used were de-identified prior to receipt by our laboratory.

### Cell culture

#### Tumor colonoid culture

Mouse tumor colonoids harboring *Apc*, *Kras*^G12D^, and *p53* mutations (AKP) and tdTomato introduced by CRISPR-Cas9 editing, were kindly provided by the Prof. Omer Yilmaz (Massachusetts Institute of Technology, USA) and cultured as previously described^39^. Briefly, AKP organoids were embedded 10 µL per dome in Cultrex Reduced Growth Factor Basement Membrane Matrix (BME; R&D Systems) and cultured in basal medium consisting of Advanced DMEM/F12 (Gibco) supplemented with 1% penicillin-streptomycin, GlutaMAX (Gibco), and 5% FBS. Tumor colonoids were passaged at a ratio of 1:3 or 1:4 and prepared for orthotopic injection as described above.

#### MAIT cell expansion from human PBMCs

MAIT cells were expanded from human PBMCs as previously described^45^. Briefly, PBMCs were cultured for 18 h with 5-A-RU (1 μg/mL) and methylglyoxal (MGO, 100 μM) as a positive control, or with βHB, NaCl, or AcAc (0.05 – 10 mM). After 18 h, culture medium [ImmunoCult-XF medium (STEMCELL Technologies) supplemented with 8% CT Immune Cell Serum Replacement (ThermoFisher Scientific) and 1% penicillin-streptomycin] was replaced with medium containing human IL-2 (33.3 ng/mL; PeproTech). IL-2-containing medium was refreshed every 3 d and cultures were maintained for up to 10 d. MAIT cell expansion, MR1 expression, and other phenotypes were quantified by flow cytometry.

#### MR1 blocking with anti-MR1 antibody

Purified anti-MR1 antibody (clone 26.5) or isotype control (BioLegend) was added at 30 μg/mL, as previously described^57^. Blocking antibody was added to cultures 30 min prior to the start of treatment (in expansion assays) or prior to combining MAIT and Caco2 cells (in co-culture experiments).

#### MAIT cell purification by magnetic separation

Following expansion, MAIT cells were stained with MR1:5-OP-RU PE tetramer (see Flow Cytometry, above) and purified using the EasySep Release Human PE Positive Selection Kit (STEMCELL Technologies) according to the manufacturer’s instructions. After bead enrichment, purity and viability were assessed by flow cytometry, and cells were used for co-culture, cytotoxicity assays, and functional analyses.

#### Caco2 cell culture

Caco2 cells (ATCC HTB-37) were maintained in Dulbecco’s Modified Eagle Medium (DMEM; Corning) supplemented with 10% fetal bovine serum and 1% penicillin-streptomycin. Cells were cultured at 37 °C with 5% CO₂ and passaged at 70-80% confluence using TrypLE Express. Only cells between passages 6 and 24 were used for experiments.

#### MAIT-Caco2 co-culture assays

Co-cultures were established when Caco2 cells reached ∼25% confluency, 18 h after seeding. Medium was removed and purified MAIT cells were added in ImmunoCult medium supplemented with 8% Immune Cell Serum Replacement at MAIT:Caco2 ratios of 16:1, 4:1, 1:1, 0:1, or 1:0. Co-cultures were maintained for 24 h and analyzed by flow cytometry. Apoptosis was assessed using Apotracker Green (BioLegend; 1:200), and viability with LIVE/DEAD Fixable Near-IR Stain (ThermoFisher Scientific). Early apoptotic cells were identified as live Apotracker^+^ cells. MAIT cell-specific killing was determined by subtracting the proportion of dead cells in Caco2-only cultures from that in Caco2-MAIT co-cultures for each treatment condition.

### LC-MS measurement of 5-A-RU, 5-OP-RU and RL-7-Me

#### Cell free 5-A-RU condensation assays

50 µL reactions containing 0.2 mM 5-A-RU and 0.6 mM metabolite reactant (MGO, Na-AcAc [prepared in-house], Li-acetoacetate [Sigma Aldrich; analytical standard], β-hydroxybutyrate, EAA, or acetone) were assembled in 10 mM HEPES buffer (pH 7.4, MS-grade water) in 1.5 mL microcentrifuge tubes and incubated at 37 °C with shaking (700 rpm, Thermo Mixer C). For time-course studies, reactions were sampled up to 3 h; otherwise, incubations proceeded for 30 min. Reactions were chilled on ice to quench, then diluted 1:5 in cold acetonitrile/water (1:1, v/v) for LC-MS analysis. Due to the instability of 5-A-RU and its condensation products, samples were analyzed by LC-MS immediately.

#### PBMC metabolite extraction

PBMCs were cultured as described above with 1 μg/mL 5-A-RU and either 50 μM MGO, 5 mM Na-AcAc, or 5 mM NaCl for 3 h. Cultures were centrifuged at 450 × g for 4 min at 4 °C. After collecting 10 μL of spent media for analysis, cell pellets were resuspended in 1 mL cold 1× dPBS and counted by Trypan Blue staining. Cells were centrifuged again (450 × g, 4 min, 4 °C), and pellets were extracted with 250 μL cold methanol/water (80:20, v/v). 10 µL of the collected spent media were extracted with 190 μL methanol/water (80:20, v/v). Samples were vortexed for 10 min at 4°C and centrifuged at maximum speed for 10 min at 4°C. The clarified extracts were transferred directly to LC-MS vials and immediately analyzed by LC-MS.

#### Liquid-chromatography mass spectrometry (LC-MS) analysis

LC-MS analysis of polar metabolites was performed using a Vanquish HPLC (Thermo Scientific) coupled to an Exploris 480 Orbitrap mass spectrometer with a heated electrospray ionization (H-ESI) source in negative-ion mode. Samples were loaded from a 4 °C autosampler onto a SeQuant ZIC-pHILIC column (2.1 × 150 mm, 5 µm; Millipore Sigma) at 25°C. Injection volumes were: 2 µL for in vitro reaction extracts, 8 µL for PBMC extracts, and 5 µL for PBMC spent media. Elution was performed as follows: 0-10 mins, linear gradient from 80% B to 50% B; 10-10.5 mins, linear gradient from 50% B to 20% B; 10.5-12.5 mins, 20% B; 12.5-13 mins, linear gradient from 20% B to 80% B; 13-20 mins, 80% B. Flow was 150 µL min⁻¹, with eluent diverted to waste from 10-20 min. H-ESI parameters: ion transfer tube 325°C, vaporizer 230°C, sheath/aux/sweep gas 35/7/1 units, negative mode spray voltage 2800 V, RF lens 60%. Single ion monitoring (SIM) was used for targeted detection of metabolite reactants and 5-A-RU-derived products (see Supplementary Table 7); SIM settings: resolution 90,000 FWHM, AGC “standard,” max injection time “auto,” scan width 2 m/z. Metabolite abundance was quantified using Xcalibur 4.1.50 (Thermo Scientific). PBMC metabolite intensities were normalized to live cell number.

### LC-MS measurement of methylglyoxal (MGO)

Li-AcAc and Na-AcAc (5 mM in PBS) were added to 8 mL glass vials and stirred using magnetic bars for 3 h at 37°C under ambient or anaerobic conditions. For time course experiments, 100 µL of samples were transferred to a 4-mL vial and treated with an equal volume of 16 mM 4-methoxy-o-phenylenediamine (4-PDA; Sigma-Aldrich) in PBS buffer for derivatization. The reaction proceeded for 30 min at rt, and an aliquot (50 µL) of the suspension was analyzed using HPLC-MS coupled with a C18 column (Phenomenex, Kinetex, EVO C18, 100 × 4.6 mm) and a gradient solvent system (10%-100% acetonitrile/water over 10 min with 0.1% formic acid; flow rate: 0.7 mL/min; UV detection: 210, 230, 254, 280, and 360 nm). The reaction product, 2-methyl-quinoxaline, was detected based on the UV spectrum and +ESI MS data ([M+H]+ m/z 175.1). Standard MGO (0.5 mM in PBS/water) was also derivatized with 4-PDA, and an aliquot (1 µL) was analyzed under the same conditions for reference. LC-MS data tables for measurement of 5-A-RU, 5-OP-RU, RL-7-Me and MGO are provided in Supplementary Data 8.

### Statistical analysis

The statistical tests used, the number of samples (n), and the number of independent experimental repeats are described in figure legends. Statistical analyses were calculated using Graphpad PRISM v10.6.0 or newer and *P* ≤ 0.05 was considered statistically significant for all experiments.

### Graphical representations

Diagrams and schemata were created using ChemDraw v25.0, Adobe Illustrator v29.8 or Biorender.

## Supporting information

Supplementary Table 1: DEGs for Total Tumor Cell Clusters

Supplementary Table 3: Mouse Flow Antibodies

Supplementary Table 4: Human Flow Antibodies

Supplementary Table 5: Human PBMC Composition

Supplementary Table 6: 10x Flex mm10 Probeset

Supplementary Table 7: Selected Ion Monitoring

## DATA AVAILABILITY

scRNA-seq datasets and LC-MS raw files are made publicly available at the time of publication.

## ACKNOWLEDGEMENTS

The authors would like to thank past and present members of the Garrett lab for their discussion of this work. We thank Prof. Omer Yilmaz (Massachusetts Institute of Technology, USA) for providing AKP tumor organoids and Prof. Mansour Haeryfar (University of Western Ontario, CA) for providing MAIT^Cast^ x *MR1^−/−^* mice. We thank the NIH Tetramer Core Facility (contract number 75N93020D00005) for providing the following tetramers: Human MR1:5-OP-RU PE, Human MR1:6FP PE, Mouse MR1:5-OP-RU PE, Mouse MR1:6FP PE, Mouse MR1:5-OP-RU BV421, Mouse MR1:6FP BV421, Mouse CD1d:PBS-57 APC, and Mouse CD1d:Unloaded control APC. We thank the Koch Institute/Whitehead Institute Metabolite Profiling Core Facility for mass spectrometry instrumentation and maintenance, Dr. Charles Vidoudez at the Harvard Center for Mass Spectrometry, Nicole Ramirez at the Bauer Core Facility, and Meghan MacDonald and Rosa Perez for colony maintenance. M.G.V.H. acknowledges support from the MIT Center for Precision Cancer Medicine, the Ludwig Center at MIT, and the NCI (R35CA242379, P30CA014051). This work was supported by a Dana-Farber Cancer Institute Cancer Immunology Training Grant, Ruth L. Kirschstein NRSA T32CA207201 to S.L.C. (2020 - 2022), and Cancer Grand Challenges C51981-A29063 to W.S.G.

## AUTHOR CONTRIBUTIONS

S.L.C. and W.S.G. conceptualized the research. S.L.C. designed and performed experiments, analyzed and visualized the data, and wrote and revised the manuscript. G.E.T. and N.A. performed orthotopic injections. E.H.R. and Y.H.S. performed LC-MS experiments. E.H.R. and S.B. performed data processing. J.N.G. analyzed histology. D.F.-P., M.M., G.N., A.F.L. assisted with experiments. S.P.C., J.C., M.G.V.H. and W.S.G. provided resources. W.S.G. supervised the project, provided funding, and reviewed and edited the manuscript.

## EXTENDED DATA FIGURES

**Extended Data Fig. 1:**
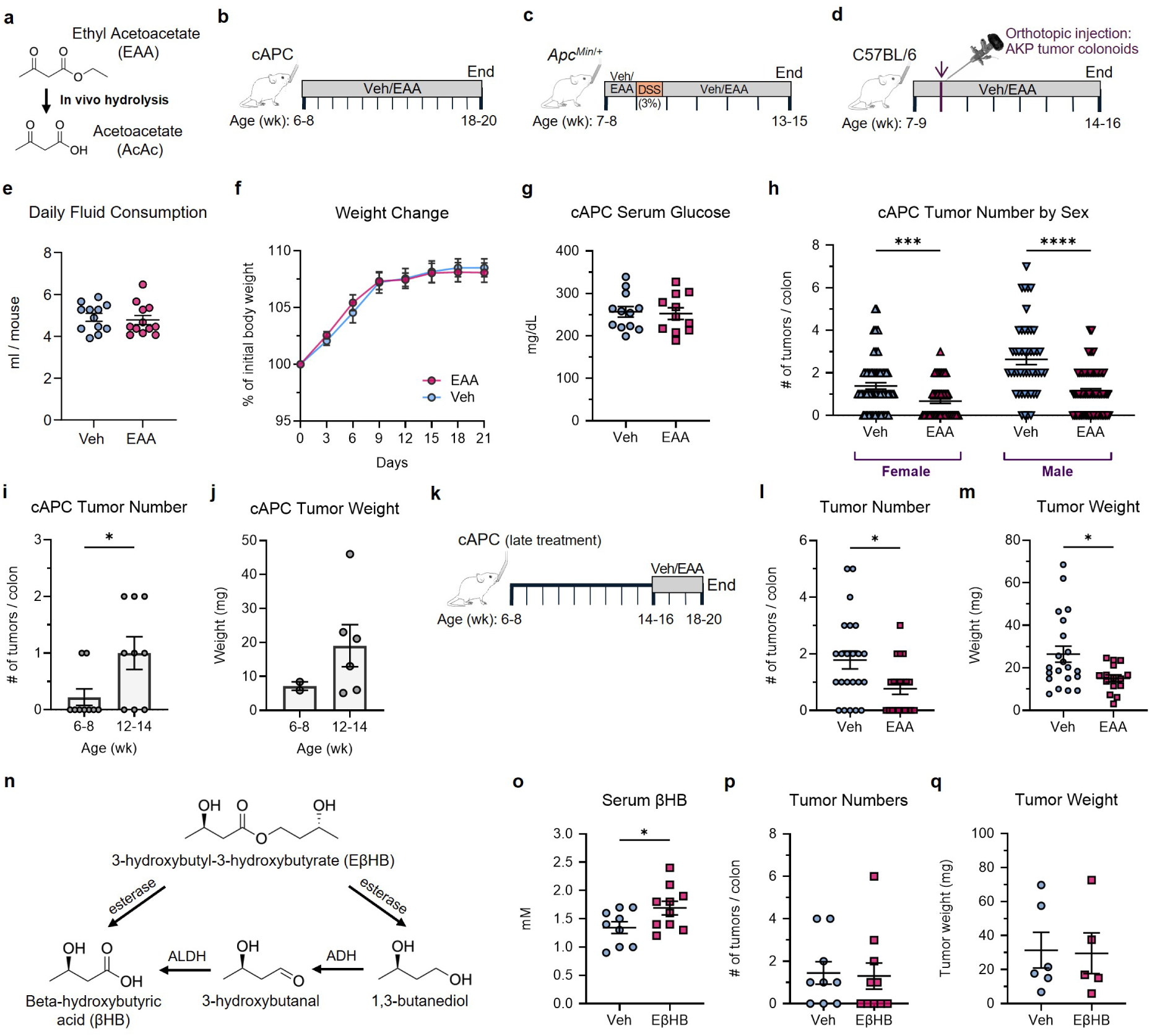
Ketone ester treatments and outcomes in mouse models of colorectal cancer. **a**, Schematic of hydrolysis of ethyl acetoacetate (EAA) to acetoacetate (AcAc). **b**-**d**, Experimental designs for *Cdx2*^Cre^ x *Apc*^flox/+^ (cAPC) (**b**), *Apc*^Min/+^ + dextran sulfate sodium (DSS) (**c**), and orthotopic AKP tumor colonoid injection (**d**) models of colorectal cancer (CRC). Vertical lines indicate weeks. **e**, Daily fluid consumption in cAPC mice administered EAA or Vehicle (Veh) control (n = 12 / group). **f**, Weight change over 21 days following initiation of treatment (n = 9-13 / group). **g**, Serum glucose levels in cAPC mice (n = 11-12 / group). **h**, Tumor number in cAPC mice following treatment and grouped by sex (n = 47-70 / group). cAPC tumor number (**i**) and weight (**j**) in untreated mice at 6-8 or 12-14 weeks of age (n = 9 mice / group). **k**, Schematic depicting time-course for treatment experiments for cAPC mice with established tumors. Tumor number (**l**) and weight (**m**) in cAPC mice following treatment (n = 15-23 / group). **n**, Pathways of esterized (R)-3-hydroxybutyl (R)-3-hydroxybutyrate (EβHB) to β-hydroxybutyrate (βHB). **o**, Serum βHB concentration in cAPC mice following treatment with EβHB or control (n = 9-10 / group). Tumor numbers (**p**) and mass (**q**) in cAPC mice following treatment with EβHB or control (n = 5-10 / group). Data from three independent experiments (**e,i,j,o**-**q**), two (**f**,**g**), four (**l**,**m**), and 17 (**h**). Lines and error bars represent mean ± s.e.m. Statistical significance was determined by Mann-Whitney *U* test (**e**,**g**,**i**-**q**) or Kruskal-Wallis test (**h**). \**P* < 0.05, \*\*\**P* < 0.001, \*\*\*\**P* < 0.0001.

**Extended Data Fig. 2:**
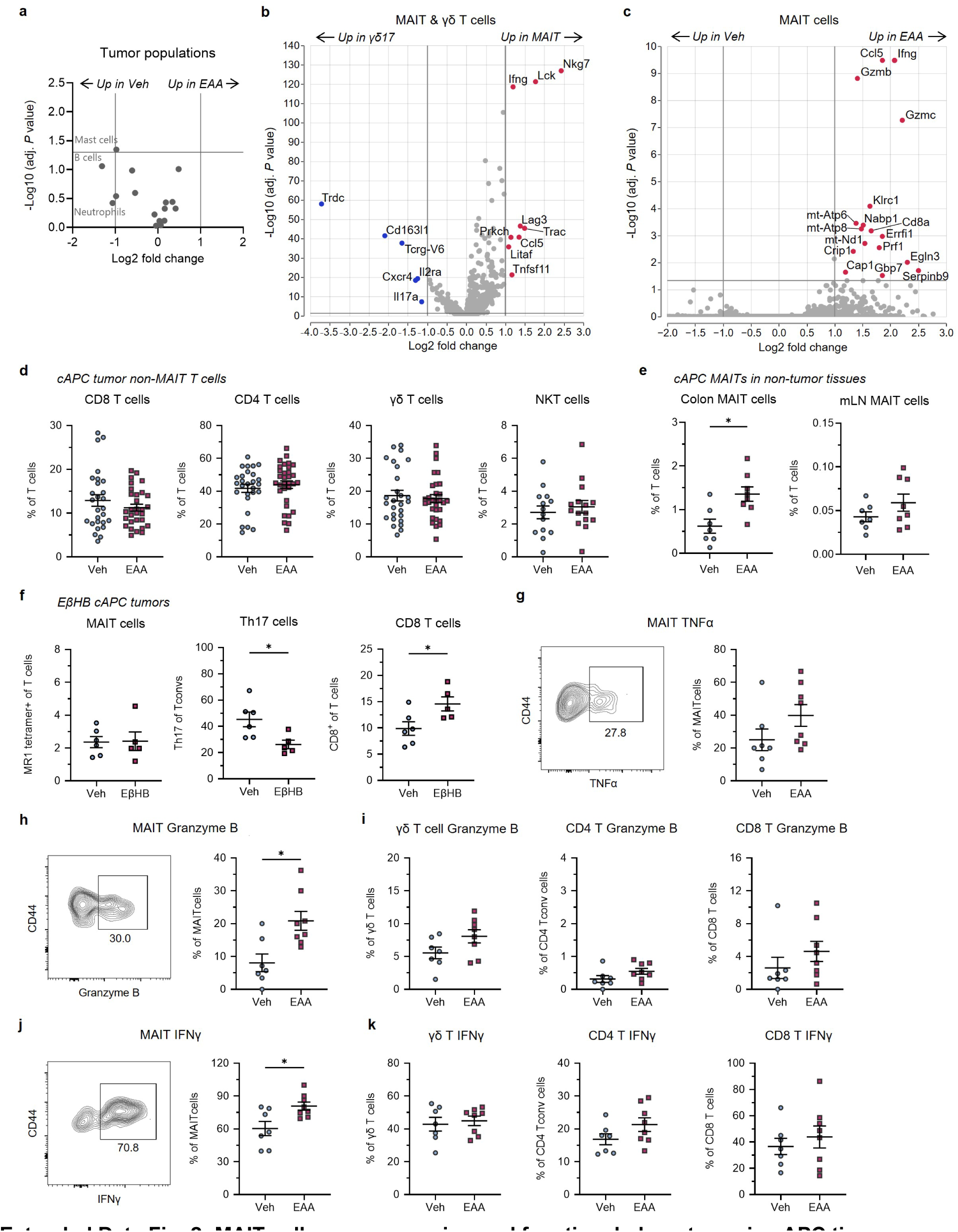
MAIT cell gene expression and functional phenotypes in cAPC tissues. **a**, Volcano plot of cAPC tumor cell populations (as shown in Fig. 2a) after treatment. Genes are labeled if Log_2_ fold change (Log_2_FC) > 1 or adjusted *P* < 0.05. **b**, Volcano plot of differentially expressed genes (DEGs) comparing MAIT and γδ17 cells, with genes labeled if Log_2_FC > 1 and adjusted *P* < 0.05. Red and blue dots indicate increased expression in MAIT or γδ17 cells, respectively. **c**, Volcano plot of MAIT cell DEGs following EAA treatment. Genes are labeled if Log_2_FC > 1 and adjusted *P* value < 0.05. Red dots indicate increased expression in EAA-treated tumors. **d**, Proportion of T cells from cAPC tumors that are CD8 T cells, CD4 T cells, γδ T cells, or CD1d tetramer^+^ natural killer T (NKT) cells (n = 15-30 / group). **e**, Proportion of T cells that are MAIT cells in adjacent colon lamina propria (LP) or mesenteric lymph nodes (mLNs) of tumor-bearing cAPC mice (n = 7-8 / group). **f**, Proportion of T cells that are MAIT, Th17, or CD8 T cells in cAPC tumors treated with EβHB or vehicle control (n = 5 / group). **g**, Representative flow cytometry plot of MAIT cell tumor necrosis factor alpha (TNFα), and the proportion of TNFα^+^ MAIT cells (n = 7-8 / group). **h**, Representative flow cytometry plot of MAIT cell granzyme B, and the proportion of granzyme B^+^ MAIT cells (n = 7-8 / group). **i**, Proportion of granzyme B^+^ γδ T cells, CD4 T cells or CD8 T cells in cAPC tumors (n = 7-8 / group). **j**, Representative flow cytometry plot of MAIT cell interferon gamma (IFNγ), and the proportion of IFNγ^+^ MAIT cells (n = 7-8 / group). **k**, Proportion of IFNγ ^+^ γδ T cells, CD4 T cells or CD8 T cells in cAPC tumors (n = 7-8 / group). Data are from six independent experiments (**d**) or three independent experiments (**e**-**k**). Lines and error bars represent mean ± s.e.m. Statistical significance was determined by Wilcoxon rank-sum test with Benjamini-Hochberg false discovery rate (FDR) correction (**a**-**c**) or Mann-Whitney U test (d-k). \**P* < 0.05.

**Extended Data Fig. 3:**
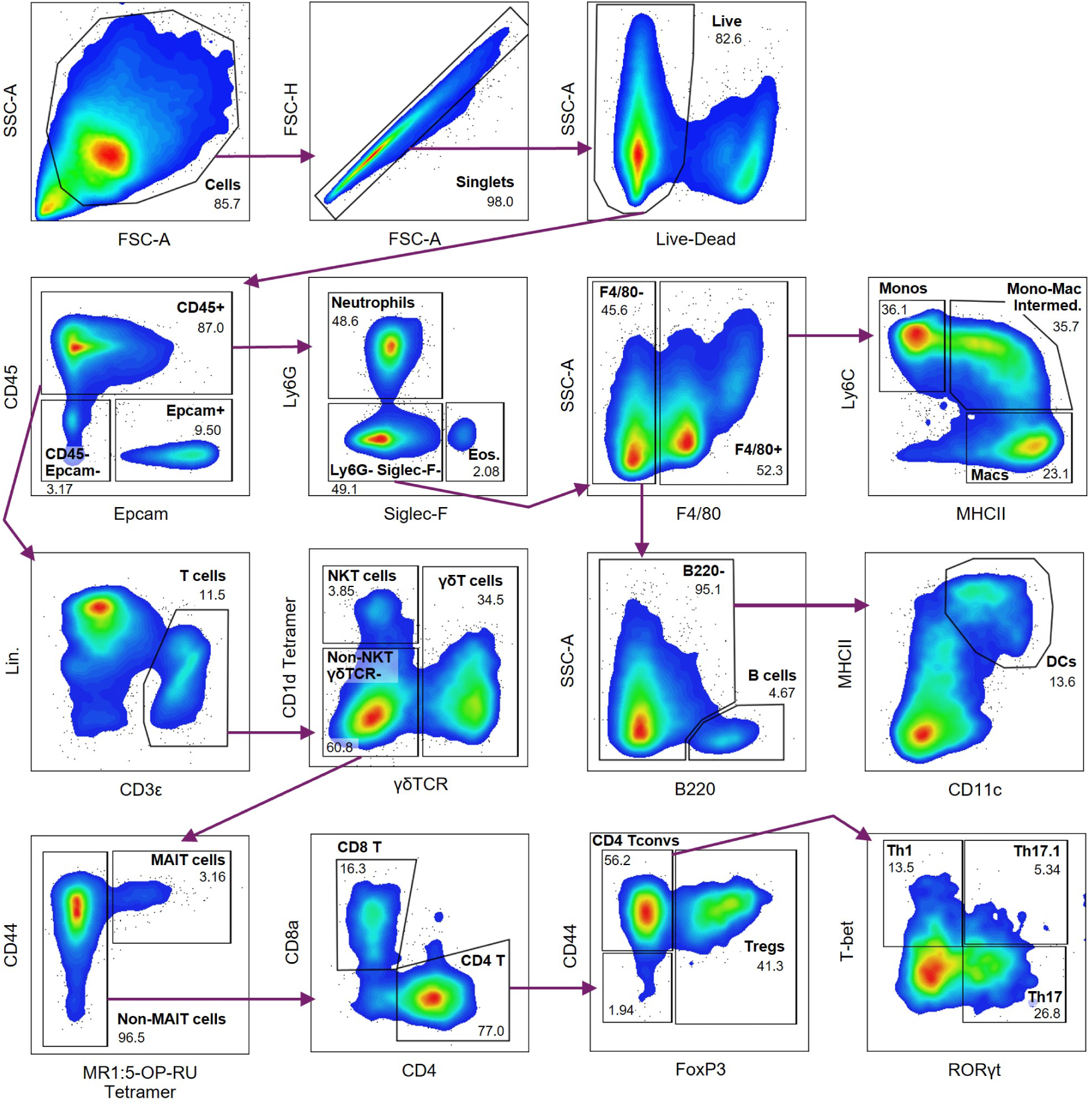
Representative gating strategies for mouse experiments. Flow cytometry gating strategy for cell populations in cAPC tumors. Population names are in bold and proportions are indicated. Arrows represent gating hierarchy.

**Extended Data Fig. 4:**
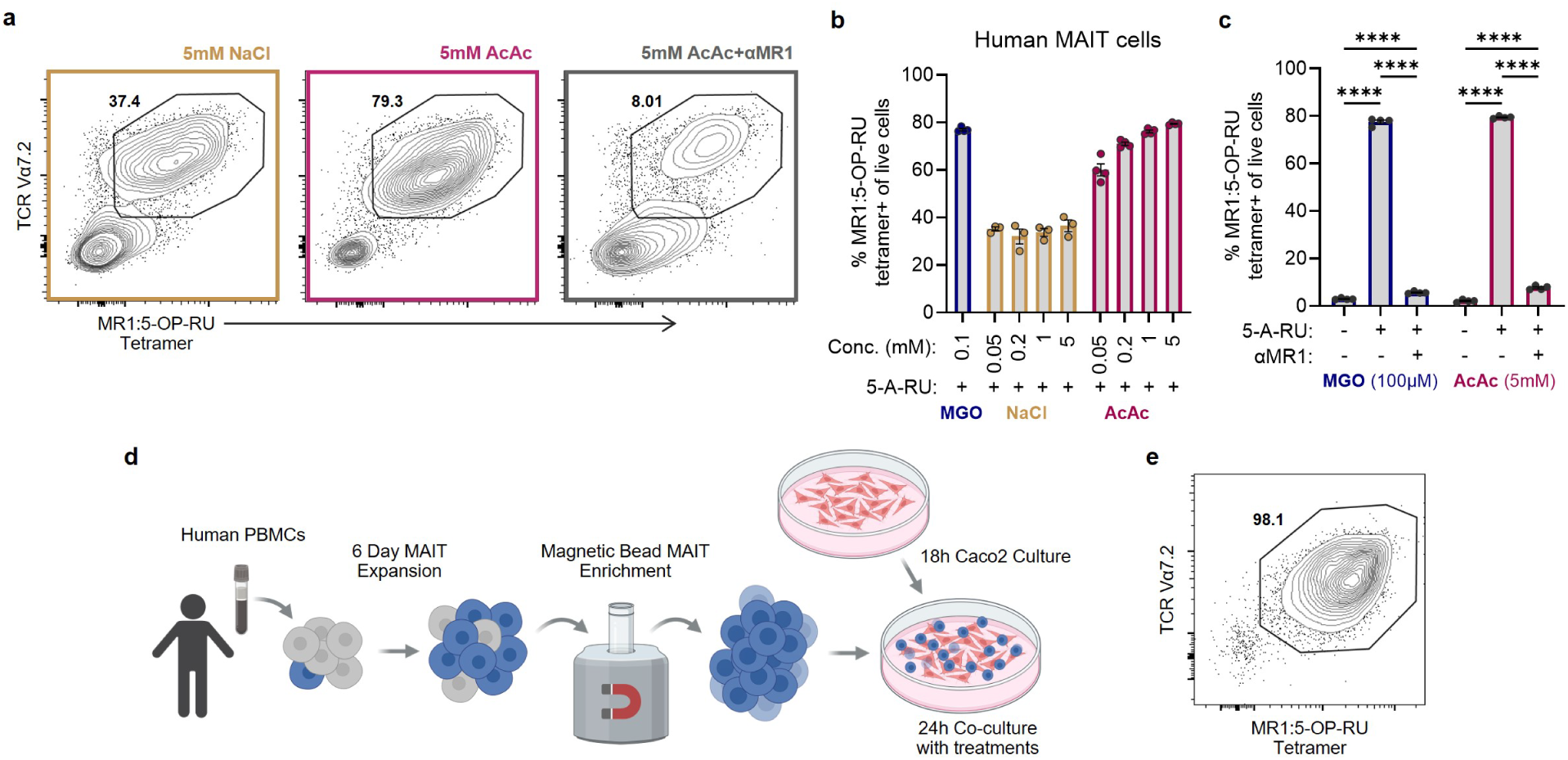
Expansion and enrichment of human MAIT cells from PBMCs. **a**, Representative flow cytometry plots of TCR Vα7.2^+^ MR1 tetramer^+^ MAIT cells following 6 d *of in vitro* expansion from whole PBMCs from a second donor (25C), with treatments indicated. **b**, Proportion of PBMCs that are MR1 tetramer^+^ MAIT cells after 6 d of culture with indicated treatments (n = 3-4 / group). **c**, Proportion of MR1 tetramer^+^ MAIT cells following culture with MGO or AcAc, with or without 5-A-RU and αMR1 antibody (n = 4 / group). **d**, Workflow for human MAIT-Caco2 co-cultures. **e**, Representative flow cytometry plot of TCR Vα7.2^+^ MR1 tetramer^+^ MAIT cells following magnetic bead enrichment. Lines and error bars represent mean ± s.e.m. Statistical significance was determined by one-way ANOVA with Tukey’s multiple comparison test. \*\*\*\**P* < 0.0001.

**Extended Data Fig. 5:**
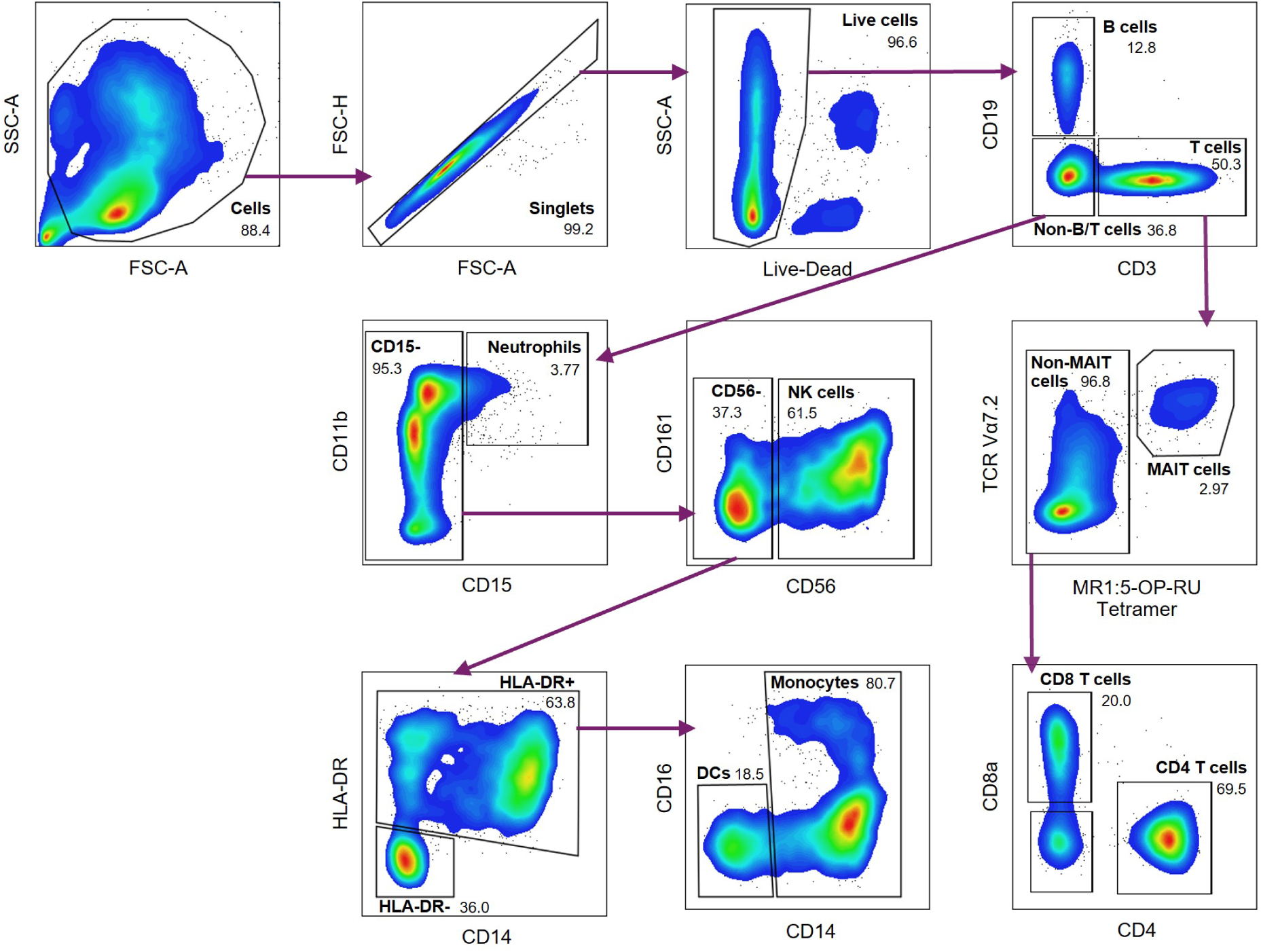
Representative gating strategy for human cells. Flow cytometry gating strategy for human PBMCs. Population names and proportions of total live cells indicated. Arrows represent gating hierarchy.

**Extended Data Fig. 6:**
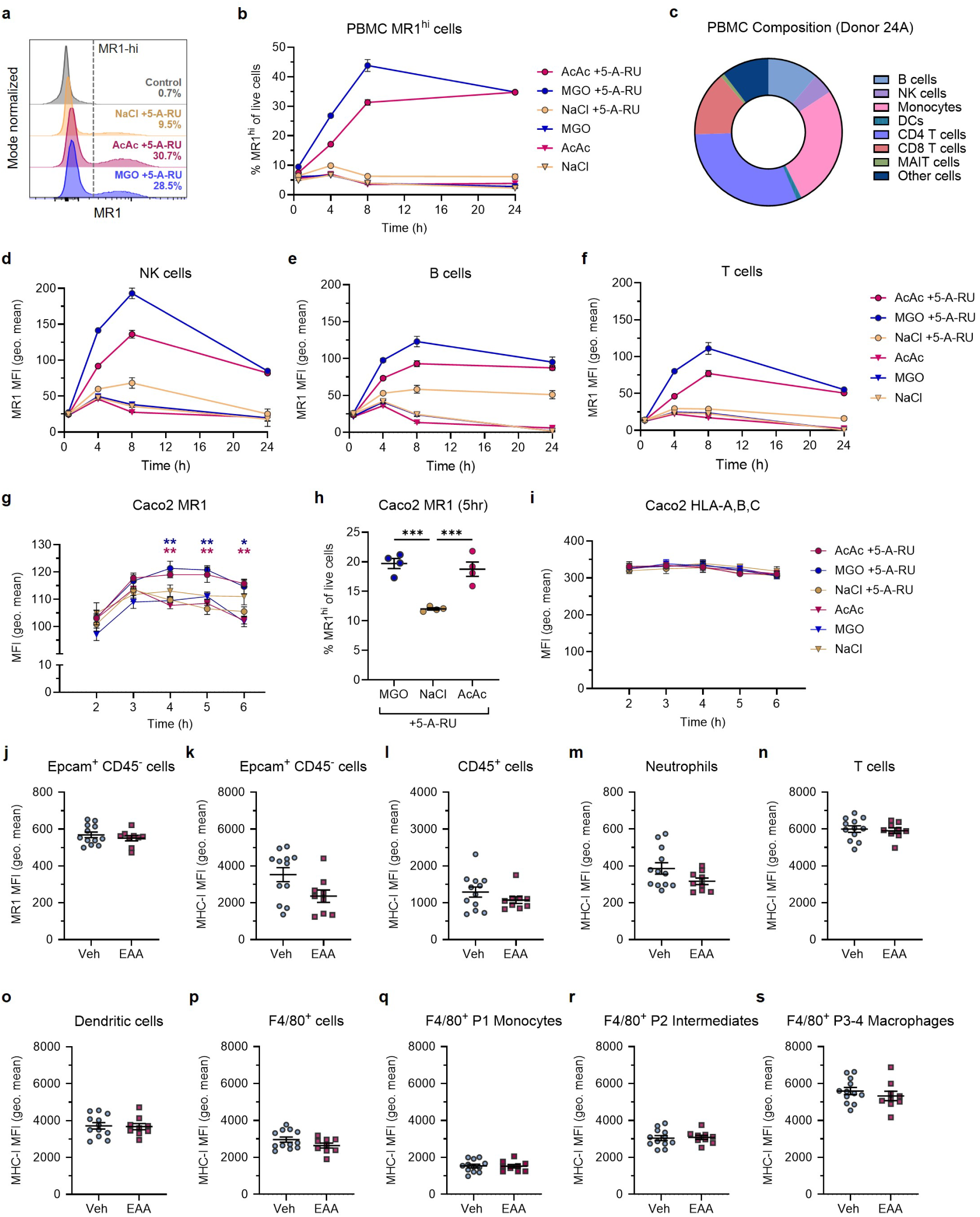
Surface MR1 expression dynamics in human and mouse cells. **a**, Representative histograms of surface MR1 geometric (geo.) MFI by total human PBMCs following 24 h of culture with treatments indicated. Dashed line indicates gating of MR1^hi^ cells. The proportion of cells that are MR1^hi^ are labeled. **b**, Time course showing the proportion of MR1^hi^ cells in human PBMCs (n = 5 / group). **c**, Composition of human PBMCs (donor 24A) determined by flow cytometry. **d**-**f**, Time course of surface MR1 geo. MFI) in human NK cells, B cells, or T cells (n = 5 / group). At 24 h, AcAc/MGO+5-A-RU vs AcAc/MGO: *P* < 0.0001 (**b**,**d,e,f**). **g**, Surface MR1 staining (geo. MFI) in Caco2 cells cultured with treatments in legend (n = 4 / group). **h**, The proportion of Caco2 cells that are MR1^hi^ after 5 h culture (n = 4 / group). **i**, Time course of HLA-A,B,C staining (geo. MFI) in Caco2 cells (n = 4 / group). **j**, Surface MR1 expression by cAPC tumor EpCAM^+^ CD45^−^ cells (n = 9-12 / group). **k**-**s**, MHC I expression by cAPC tumor populations (n = 9-12 / group). Data are representative of two independent experiments (**a**-**i**) or from three independent experiments (**j**-**s**). Lines and error bars represent mean s.e.m. (**g**-**s**). Statistical significance was determined by two-way ANOVA with Geisser-Greenhouse correction and Tukey’s multiple comparison test (**b**,**d,e,f**), one-way ANOVA with Tukey’s multiple comparison test (**g**-**h**) or Mann-Whitney *U* test (**j**-**s**). \**P* < 0.05, \*\**P* < .01, \*\*\**P* < 0.001.

**Extended Data Fig. 7:**
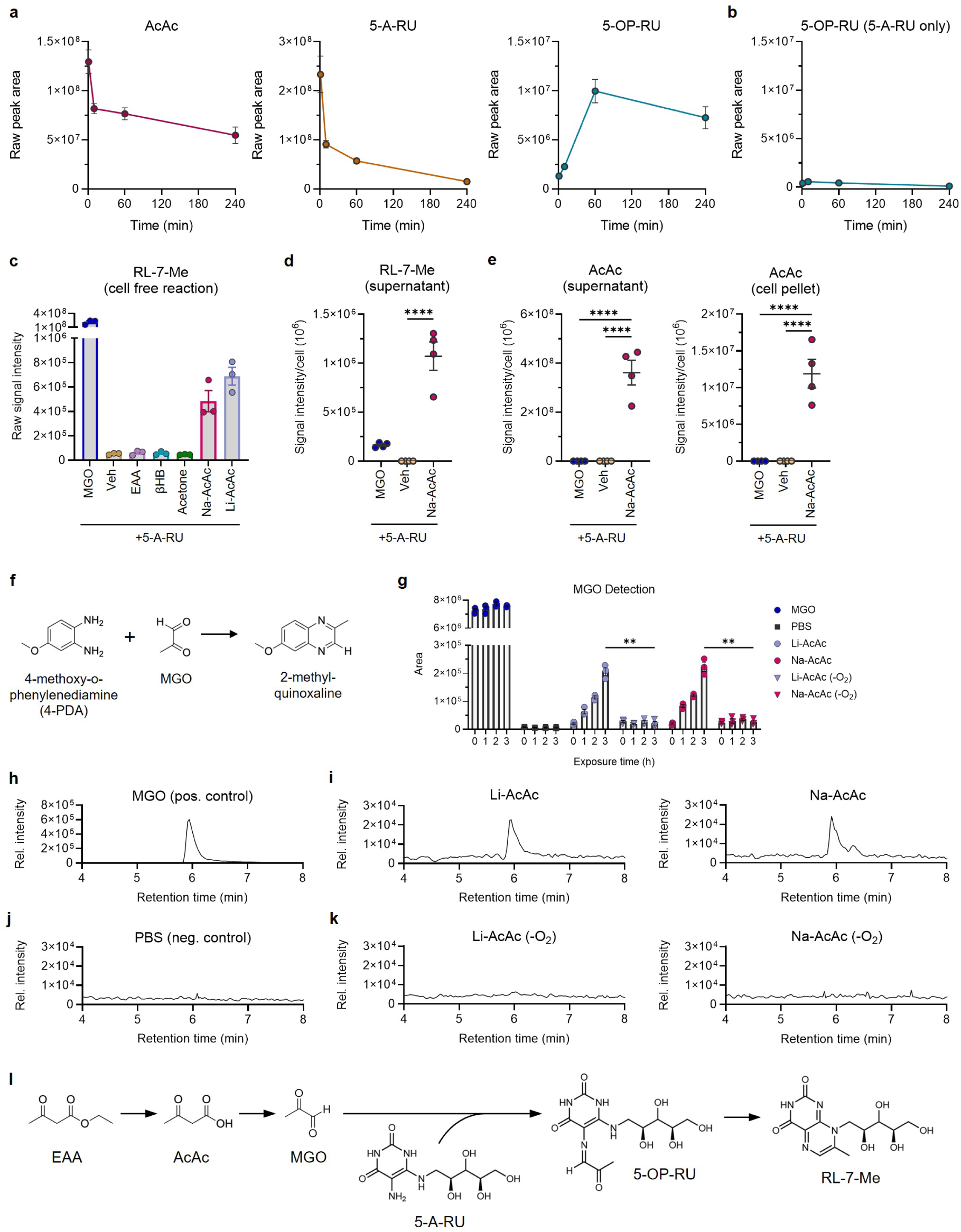
Acetoacetate forms 5-OP-RU via methylglyoxal. **a**, LC-MS measurement of AcAc and 5-A-RU consumption and 5-OP-RU generation during 240 min incubation of AcAc and 5-A-RU (n = 3 / group). **b**, Measurement of 5-OP-RU in 5-A-RU only control samples (n = 3 / group). **c**, 7-methyl-8-D-ribityllumazine (RL-7-Me) measured by LC-MS following 30 min reactions with 5-A-RU plus MGO, Veh, EAA, βHB, acetone, Na-AcAc, or Li-AcAc (n = 3 / group). **d**, RL-7-Me in supernatants (left) or cell pellets (right) after 3 h PBMC culture (n = 4 / group). **e**, AcAc in supernatants (left) or cell pellets (right) after 3 h PBMC culture (n = 4 / group). **f**, Schematic of MGO derivatization with 4-methoxy-o-phenylenediamine (4-PDA) to produce 2-methyl-quinoxaline, which is detected by LC-MS. **g**, Measurement of MGO showing individual data points and MGO reference samples (n = 3 / group). **h**, Representative extracted ion chromatogram (EIC) of MGO detection for MGO positive control sample at 3 h. **i**, Representative EICs of MGO detection for Li-AcAc and Na-AcAc samples at 3 h. **j**, Representative MGO EICs for PBS-only negative control sample, and **k**, Representative EICs for Li-AcAc and Na-AcAc under anaerobic (-O_2_) conditions at 3 h. **l**, Diagram of EAA conversion to AcAc, subsequent degradation to MGO, adduct formation with 5-A-RU to generate 5-OP-RU, and formation of the lumazine degradation product RL-7-Me. Data are representative of two independent experiments (**a,b,d,e**) or three independent experiments (**c**). Lines and error bars represent mean ± s.e.m. (**a**-**g**). Statistical significance was determined by one-way ANOVA with Tukey’s multiple comparison test (**d,e**) or two-way ANOVA with Geisser-Greenhouse correction and Tukey’s multiple comparison test. \**P* < 0.05, \*\**P* < .01, \*\*\*\**P* < 0.0001.

## SUPPLEMENTARY TABLES

**Supplementary Table 1: Top DEGs for Clusters of Total Tumor Cells**

**Supplementary Table 2: Top DEGS for Clusters of Tumor Lymphocytes**

**Supplementary Table 3: Mouse Flow Antibodies**

**Supplementary Table 4: Human Flow Antibodies**

**Supplementary Table 5: Human PBMC Composition**

**Supplementary Table 6: 10x Flex mm10 Probeset**

**Supplementary Table 7: Selected Ion Monitoring**

**Supplementary Table 8: LC-MS Tables**

